# ProMPt: A modular preclinical platform for functional modelling of prostate cancer heterogeneity and therapeutic vulnerabilities

**DOI:** 10.1101/2025.11.16.688689

**Authors:** Nicole Pandell, Jacob Househam, Matteo Tartagni, Archana Thankamony, Florian Gabel, Roberto Rota, Elizabeth Flittner, Bora Gurel, Ines Figueiredo, George Seed, Antje J. Neeb, Mary Chol, Wei Yuan, John G. Clohessy, Christopher J. Tape, Adam Sharp, Johann S. de Bono, Marco Bezzi

## Abstract

Prostate cancer progression is driven by heterogenous genetic, phenotypic, and microenvironmental programs that remain challenging to model experimentally. Existing systems such as genetically engineered mouse models, xenografts, and patient-derived organoids have each advanced mechanistic insight but are limited by genetic scope, scalability, or lack of immune context. To overcome these constraints we developed ProMPt, a genetically-defined syngeneic mouse modelling platform that captures combinations of the most recurrent clinical prostate cancer genomic alterations to enable scalable *in vitro* and *in vivo* interrogation of prostate cancer evolution. Tumours derived from ProMPt organoids recapitulate the histologic and molecular diversity of human disease. Cross - species transcriptomic integration and multivariate single-cell analysis under defined culture permutations revealed conserved phenoscapes, highlighting a central role for *MYC* in disease progression and therapy resistance. Guided by these insights, preclinical intervention studies demonstrated that combined MAPK inhibition and blockade of protein translation synergistically suppressed tumour growth in castration-resistant models. This combination not only suppressed proliferation but also remodelled the tumour immune landscape, underscoring its dual epithelial and microenvironmental effects. Together, these findings establish ProMPt as a versatile framework for linking genotype, lineage plasticity, and therapeutic vulnerability in prostate cancer.

## INTRODUCTION

Prostate cancer (PC) is the most frequently diagnosed malignancy in men and a leading cause of cancer mortality worldwide(1). Despite initial dependence on androgen-receptor (AR) signalling, most tumours progress to castration-resistant prostate cancer (CRPC), a lethal stage marked by molecular, phenotypic, and microenvironmental heterogeneity(2). Genomic profiling of advanced disease has revealed intricate combinations of canonical drivers such as *PTEN* loss, *TP53* mutations, *TMPRSS2–ERG* fusion, and *MYC* amplification, interwoven with a long tail of lower-frequency alterations that remodel signalling networks and reprogram tumour–microenvironment interactions(3). These alterations converge into three major phenotypes: AR-positive (ARPC), double-negative (DNPC), and neuroendocrine (NEPC), whose relationships and evolutionary trajectories remain poorly defined(4). Loss of *TP53* and *RB1* is strongly associated with neuroendocrine differentiation, yet complete lineage conversion appears to require additional cooperating oncogenic, epigenetic, and microenvironmental cues that are still being elucidated(5,6). In contrast, the DNPC spectrum spans diverse molecular and phenotypic states, often linked to biallelic *PTEN* loss(7), Glucocorticoid receptor activity(8), and activation of WNT and FGFR/MEK signalling pathways that regulate stem-like and epithelial–mesenchymal transition (EMT) programmes whose interplay remains poorly characterised(9–12) .

This complex phenoscape underscores the need for scalable models that resolve how genetic context, lineage states, and microenvironmental cues interact to drive progression and therapy resistance, enabling both mechanistic insight and preclinical testing across the breadth of CRPC diversity. Existing systems capture aspects of prostate-cancer biology but remain limited. Human prostate cancer cell lines are few and fail to reflect the molecular and phenotypic diversity of advanced disease. Patient-derived xenografts (PDXs) and organoid cultures offer greater fidelity to patient tumours yet remain difficult to establish, are genetically heterogeneous, and cannot be easily studied within an intact immune microenvironment(13–15). Genetically engineered mouse models (GEMMs) have yielded key insights into tumour initiation and progression but are time-consuming, poorly scalable, and limited in reproducing complex multi-driver genotypes. Comparative studies in syngeneic, immunocompetent systems have therefore begun to emerge(6,16,17), providing an important foundation and highlighting the opportunity for next-generation, modular *in vivo* platforms to interrogate cancer evolution and therapeutic response.

Here, we present ProMPt, a next-generation Prostate cancer Modelling Platform based on a murine organoid biobank derived from epithelial cells engineered with clinically relevant combinations of driver alterations (*PTEN*, *TP53*, *RB1*, *MYC*, *TMPRSS2–ERG*, and *APC*)(3). Focusing on luminal-derived lines, we demonstrate active Ar signalling and show how specific genotypes modulate responses to androgen deprivation. Upon transplantation, ProMPt organoids generate CRPC tumours spanning distinct epithelial and microenvironmental states, with MYC emerging as a key determinant in *Pten*-null contexts. Integration of transcriptomic profiling with cross-species Human-Mouse (HuMo) clustering(18) revealed that ProMPt tumours mirror conserved molecular patterns of human CRPC. Systematic modulation of androgen and paracrine cues, coupled with single-cell CyTOF analysis(19), identified context-specific vulnerabilities, including dependency on EGF signalling that is bypassed by organoids with *Pten*–*MYC* co-alteration. These aggressive models exhibited intrinsically elevated protein-translation pathways, prompting *in vivo* evaluation of combined MEK and translation inhibition, which produced marked tumour suppression and remodelling of the tumour microenvironment.

Together, ProMPt provides a modular, genetically defined, and immunocompetent framework that integrates genetic diversity with scalability and flexibility for future expansion. This resource enables discovery of genotype-specific vulnerabilities, supports rational therapeutic design, and advances mechanistic understanding of prostate-cancer evolution.

## RESULTS

### A modular syngeneic organoid pipeline enables systematic modelling of prostate cancer drivers

Both luminal and basal prostate epithelial cells can initiate tumorigenesis, yet their behaviour varies with oncogenic context, and when cultured as organoids they display distinct proliferative and differentiation dynamics(20–22). In GEMMs, tumour onset and phenotype depend on the underlying genetic alterations and organoids derived from these lesions often show mixed or altered lineage marker expression, complicating direct genotype–phenotype analysis(23). To enable systematic cross-comparison, we developed the ProMPt platform based on conditional alleles and *ex vivo* induction of genomic alterations in lineage-defined, wild type-like epithelial cells. We selected drivers that combined clinical relevance with technical feasibility for syngeneic modelling. *PTEN* (Pt), *TP53* (P), *TMPRSS2–ERG* (E), *MYC* (M), *RB1* (R), and *APC* (A) are among the most recurrently altered genes in prostate cancer and are strongly implicated in progression, therapy resistance, and lineage plasticity(24–26). Each allele was available on a C57BL/6 background, ensuring compatibility with immune-competent transplantation. Analysis of large patient cohorts, including TCGA, SU2C, and MSKCC datasets, confirmed that these alterations collectively, occur in the majority of human prostate cancers, increasing from ∼60% in primary tumours to >77% in advanced and castration-resistant disease, where they exhibit complex co-occurrence patterns (Fig. 1A; Supp. Fig. 1A).

**Figure 1.**
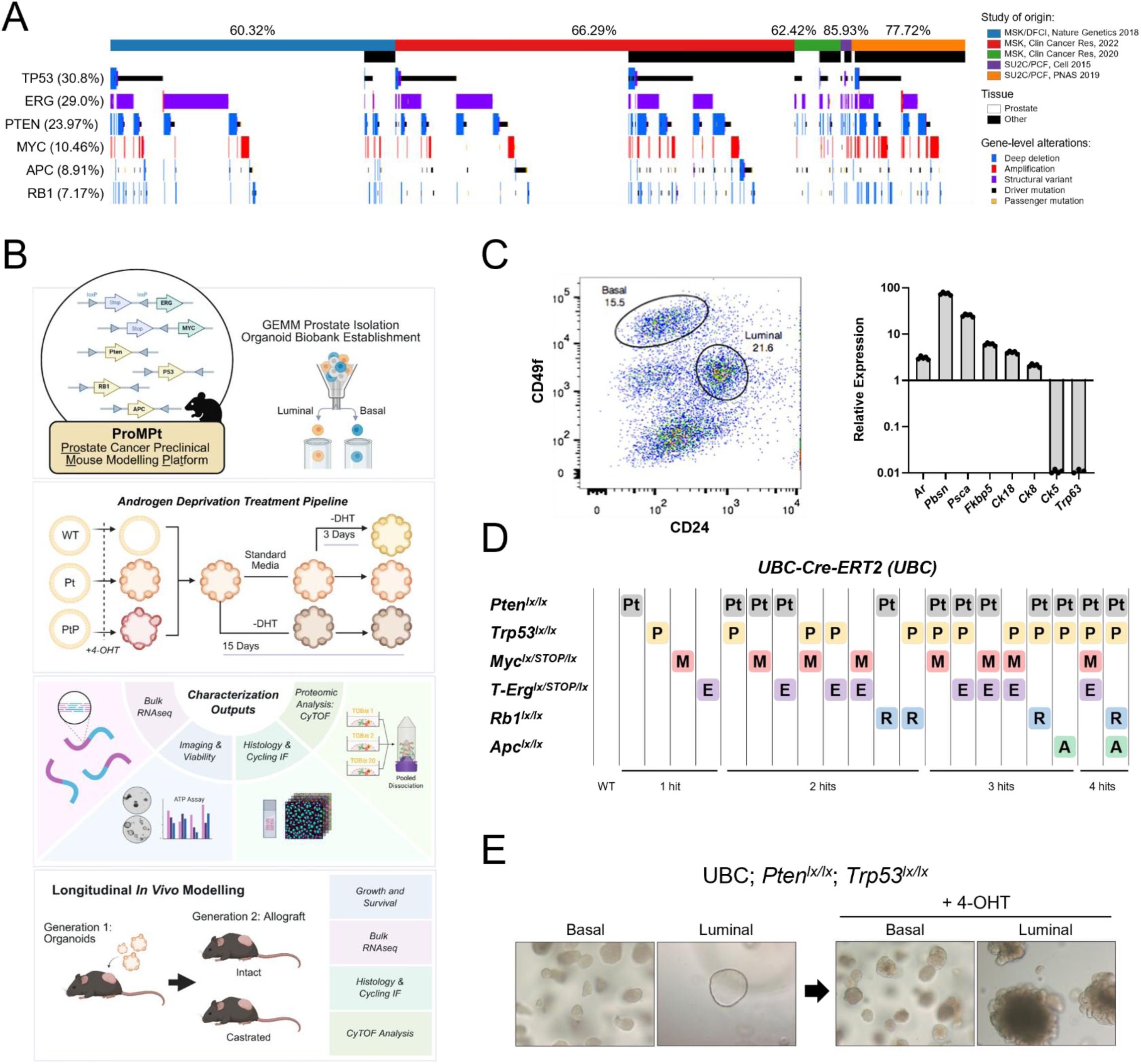
ProMPt: an organoid pipeline to model frequent prostate cancer drivers. (A) Oncoprint summarising the combined frequency of genetic aberrations across studies. Combined aberration frequency for all listed genes is shown per dataset (top) and for each specified gene across datasets (left) of primary and metastatic prostate cancer cohorts. See also Figure S1A. (B) Schematic illustrating the generation of the Prostate Cancer Preclinical Mouse Modelling Platform (ProMPt). The workflow outlines the establishment and multidimensional characterisation of ProMPt organoids under androgen deprivation and following transplantation into immunocompetent hosts. (C) Representative sorting gating of basal (Cd49f⁺Cd24⁻) and luminal (Cd49f^low^Cd24⁺) murine prostate epithelial cells (left), and quantitative RT–PCR analysis of lineage-specific transcripts in luminal cells, normalised to basal cells and Tbp expression. Each dot represents a technical replicate (n = 5). (D) Table summarising the 21 genetic profiles comprising the ProMPt platform, generated through sequential breeding of the six indicated mouse strains with each other and with the Ubc -CreERT2 line. (E) Representative brightfield images of basal and luminal organoid populations before (left) and after (right) Cre-mediated genetic recombination induced by 10-day treatment with 4-hydroxytamoxifen (4-OHT).

To introduce recombination *ex vivo* while maintaining line synchrony and avoiding viral transduction or prolonged culture, donor mice carried a Ubc-CreERT2 allele that enabled tamoxifen-inducible activation of floxed alleles. Prostates from healthy three-month-old mice of the desired genotype were harvested, and epithelial cells isolated for organoid derivation (Fig. 1B). A key feature of our approach is the independent isolation of basal (Cd49f^+^ Cd24^-^ and luminal (Cd49f^low^ Cd24^+^) cells, followed by parallel culture of each population(27). Lineage fidelity was routinely confirmed at the transcript level, with basal cells enriched for *Trp63* and *Krt5*, whereas luminal cells expressed canonical luminal markers and Ar target genes (Fig. 1C). This strategy generated two genetically defined biobanks, basal and luminal, that can be interrogated independently or directly compared to study cell-of-origin effects.

Using this framework, we established a panel of organoid lines encompassing single and combinatorial alterations across the six selected drivers (Fig. 1D), with at least three independent lines derived for each genotype. Recombination was induced immediately after cell isolation by short-term exposure to 4-hydroxytamoxifen (4-OHT) for two passages (∼10 days), enabling efficient activation or deletion of targeted alleles while preserving early-passage integrity and reducing culture-associated stress and selective pressure. Upon CreERT2 induction, both basal- and luminal-derived organoids exhibited pronounced changes, characterized by increased cellularity and morphological transformation (Fig. 1E). Molecular validation confirmed successful recombination, with expected transcript changes by qPCR and corresponding protein loss or activation by immunoblotting (Supp. Fig. 1B–D). Validated organoids were cryopreserved at low passage to establish a stable biobank, minimising lineage drift and ensuring reproducible comparisons across genotypes.

Together, these elements define the core of the ProMPt pipeline: a unified framework that integrates clinically relevant driver selection, a C57BL/6 genetic background, and inducible *ex vivo* recombination to achieve precise and reproducible modelling of prostate cancer. Through the establishment of parallel basal and luminal biobanks, and early cryopreservation to maintain fidelity, the platform offers a scalable and versatile resource for the systematic dissection and functional analysis of key genetic alterations across epithelial lineages, both *in vitro* and *in vivo*.

### Morphological diversity and transcriptional responses to androgen withdrawal in ProMPt organoids

Following the establishment of the basal and luminal ProMPt organoid biobanks, we next examined how genetic context influences organoid morphology and viability under androgen deprivation therapy (ADT). Given the well-established role of luminal epithelium as a cell of origin in prostate cancer (28–30), and the limited availability of models that retain AR signalling, luminal–derived organoids were selected as the focus for all subsequent studies. Organoids displayed a broad spectrum of morphologies, ranging from small cystic structures to multi-layered and irregular architectures. Immunostaining for luminal markers (Ck8, Trop2)(31) and basal markers (Ck5, Cd104)(32) confirmed that transformed luminal cells retained the ability to form organoids with basal–luminal organization (Fig. 2A; Supp. Fig. 2A-B). Genotype contributed markedly to this architectural diversity: organoids with *Pten* and *Trp53* double knockout backgrounds (PtP) typically developed into large, multilayered structures, whereas *MYC*-overexpressing (M) organoids exhibited dense, hyperplastic morphologies often lacking a defined lumen. Organoids with more aggressive genotypes showed loss of polarity and presence of intermediate cells. These features mirrored those observed in organoids derived from GEMMs, including multilineage differentiation and the formation of polarized epithelia(28).

**Figure 2.**
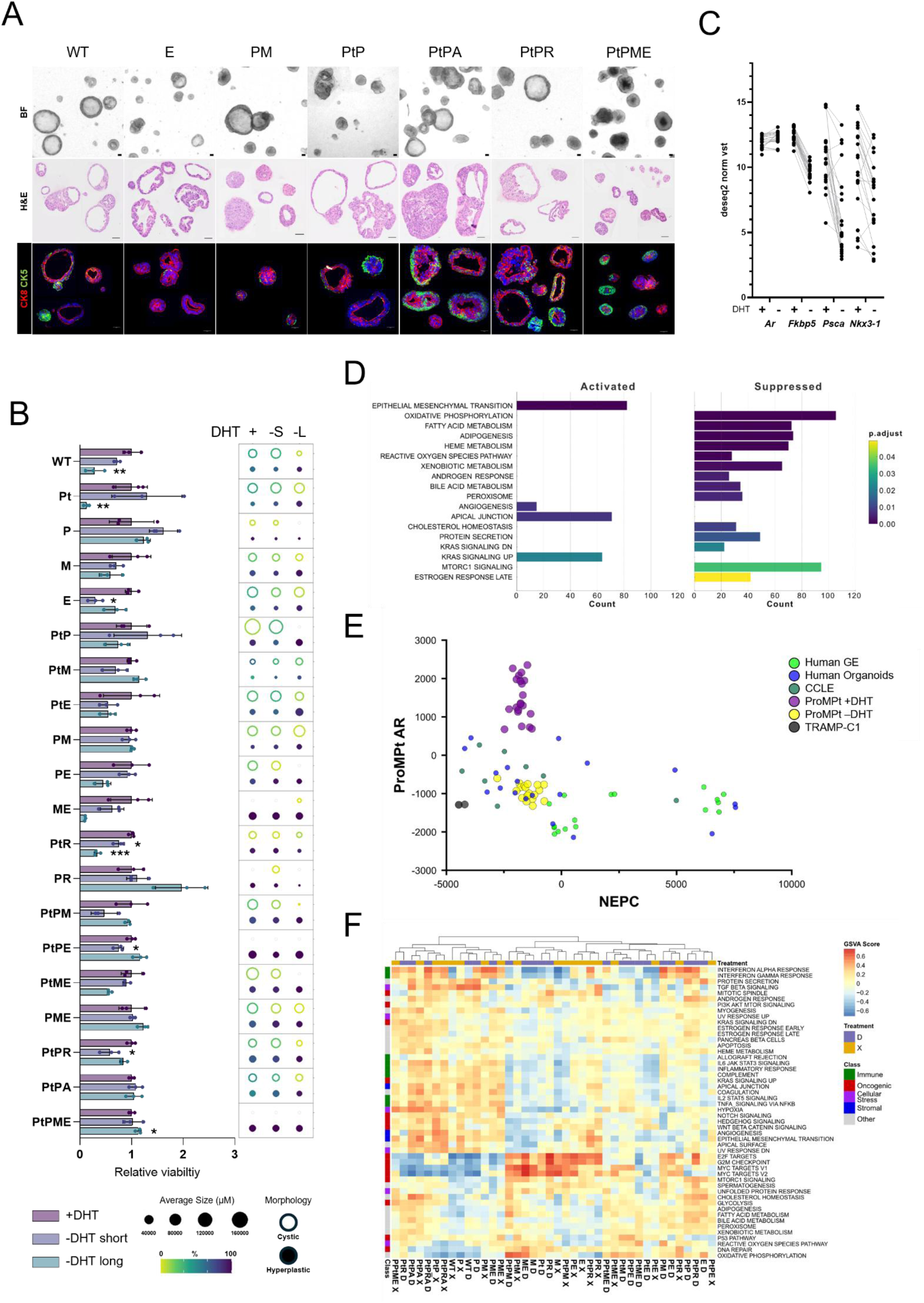
Characterization of ProMPt organoids following androgen withdrawal. (A) Representative brightfield (BF), H&E, and immunofluorescence (IF) images of ProMPt organoids following genetic recombination. Scale bars: 2 mm (BF) and 50 μm (H&E, IF). Additional organoid images and IF markers are shown in Figure S2A–B. (B) Viability (left) and morphological (right) analysis of ProMPt organoids cultured in complete medium, short-term (72 h; –S) or long-term (15 d; –L) DHT withdrawal. Viability data are normalised to the untreated condition for each genotype. Error bars indicate mean ± SD (n = 3). P values were determined using Welch’s t-test; only significant comparisons are shown (\**P* < 0.05, \*\**P* < 0.01, \*\*\**P* < 0.001). Organoid morphology (cystic versus hyperplastic) and size data are presented as a bubble plot with mean values (n = 3). (C) Normalised RNA expression levels from bulk RNA-seq of Ar, Fkbp5, Psca, and Nkx3.1 in paired +DHT and long-term DHT-withdrawn conditions. Each dot represents one ProMPt organoid model (n = 17). (D) Gene set variation analysis (GSVA) showing the top Hallmark pathways differentially enriched between untreated and long-term DHT-withdrawn organoids, ordered by significance. P values were adjusted for multiple testing. (E) Scatterplot showing expression of ProMPt-derived androgen receptor (Ar) and NEPC (10) signatures across ProMPt organoids and established prostate cancer cell lines and organoids. See also accompanying details in Supp. Table 2 and 3. (F) Unsupervised clustering of Hallmark GSVA enrichment scores. Normalised GSVA scores are scaled by row and adjusted for multiple testing correction. Pathway classes and sample treatments (D: +DHT, X: long-term –DHT) are colour coded at left and top, respectively.

To model ADT, organoids were cultured in parallel under complete medium containing dihydrotestosterone (DHT), short-term DHT withdrawal (72 h), and long-term deprivation over three passages (15 days). At endpoint, samples from all conditions were analysed by viability assays and brightfield imaging (Fig. 1B). In line with previous studies, removal of DHT produced only modest effects on proliferation across most transformed organoid lines(5,28,33) (Fig. 2B), but cultures progressively shifted toward hyperplastic morphologies, with a corresponding loss of cystic organoids. Bulk RNA sequencing confirmed robust Ar pathway activity in all lines and its consistent suppression following prolonged androgen withdrawal, marked by downregulation of canonical targets including *Fkbp5*, *Psca*, and *Nkx3.1* (Fig. 2C). Expression analysis of luminal and basal gene signatures revealed a clear reduction in luminal identity (Supp. Fig. 2C), consistent with enhanced lineage plasticity under ADT as previously reported(5).

Comparative transcriptomic analysis of all lines cultured in +DHT versus long –DHT conditions revealed extensive pathway reprogramming. Gene set enrichment analysis (GSEA) showed suppression of androgen signalling together with activation of EMT- and KRAS-associated pathways (Fig. 2D). Differential expression analysis further identified *Edil3* and *Rgs5* as upregulated under androgen deprivation, genes implicated in invasive, angiogenic, and stress-response processes(34,35), while canonical Ar targets including *Fkbp5*, *Zbtb16*, *Mmp7*, and *Nkx3.1* were among the most downregulated (Supp. Fig. 2D). From these data, we derived a 30-gene ProMPt-specific Ar signature that correlated significantly with the human Hallmark AR score in cultured lines and patient data, demonstrating cross-species concordance and improved detection of androgen-deprivation responses in murine lines (Supp. Fig. 2E-F). To assess how ProMPt models relate to established prostate cancer subtypes, we next compared their transcriptomes with reference datasets of representative prostate cancer lines. Using the Han NEPC signature(11) in combination with the ProMPt-Ar score, +DHT organoids clustered with AR-proficient prostate cancer models, whereas prolonged androgen deprivation shifted them toward the double-negative state (Fig. 2E). No evidence of neuroendocrine transdifferentiation was detected, consistent with previous reports that additional cues may be required to induce this fate *in vitro*(5). Projection against the Han mesenchymal stem–like prostate cancer (MSPC)(11) and EMT signatures further showed that ProMPt lines were enriched for MSPC programs but remained distinct from other models such the SV40 large T-cell antigen driven, and dysfunctional Rb1 and Trp53, TRAMP-C1 (Supp. Fig. 2G). Thus, androgen-deprived ProMPt organoids occupy a stem-like, lineage-plastic state without progressing to neuroendocrine or fully mesenchymal phenotypes. Finally, GSVA-based clustering of pathway enrichment profiles demonstrated that genotype was the dominant determinant of transcriptional identity (Fig. 2F). Lineage- and oncogene-associated programs segregated according to genomic background, with *MYC*-overexpressing lines forming a distinct cluster enriched for MYC and E2F target signatures. EMT-related pathways were preferentially elevated in defined subsets, suggesting that genetic context not only defines baseline epithelial identity but also modulates adaptive responses to androgen withdrawal.

Together, these results establish that ProMPt organoids recapitulate the structural and lineage heterogeneity of murine prostate models and display genotype-specific transcriptional responses to androgen deprivation that align with human CRPC phenotypes, providing a scalable platform for mechanistic and translational investigation.

### ProMPt tumours model frequent and rare CRPC subtypes

To evaluate the tumorigenic potential of ProMPt organoids *in vivo*, we established a two-step transplantation system (Fig. 3A). A subset of dissociated organoids was implanted subcutaneously into C57BL/6 mice (Gen 1), and tumours that developed were reimplanted as allografts into secondary recipients (Gen 2). Parallel cohorts of Gen 2 mice were castrated to assess androgen dependence. Tumour penetrance varied by genotype (Fig. 3B), with single-hit organoids (P and M) failing to engraft, whereas multi-hit combinations produced tumours at frequencies that increased with the number of engineered alterations. Growth kinetics further resolved three characteristic patterns (Fig. 3C; Supp. Fig. 3A–B). PtPM and PtPME organoids engrafted rapidly and grew aggressively, while triple- and quadruple-hit genotypes lacking *MYC* overexpression displayed longer and more variable latencies, suggesting an adaptation phase before outgrowth. Two-hit tumours exhibited the longest latency. Kaplan–Meier analyses mirrored these trends, with PtPM-driven tumours showing the most rapid disease progression and shortest survival, while non-*MYC* models demonstrated greater heterogeneity. Once established, all genotypes grew efficiently as Gen 2 allografts with high penetrance and short latency. Castration induced AR cytoplasmic relocalisation and frequent downregulation, yet had no significant effect on growth (Supp. Fig. 3C), indicating that these tumours represent *de novo* castration-resistant prostate cancers (CRPCs).

**Figure 3.**
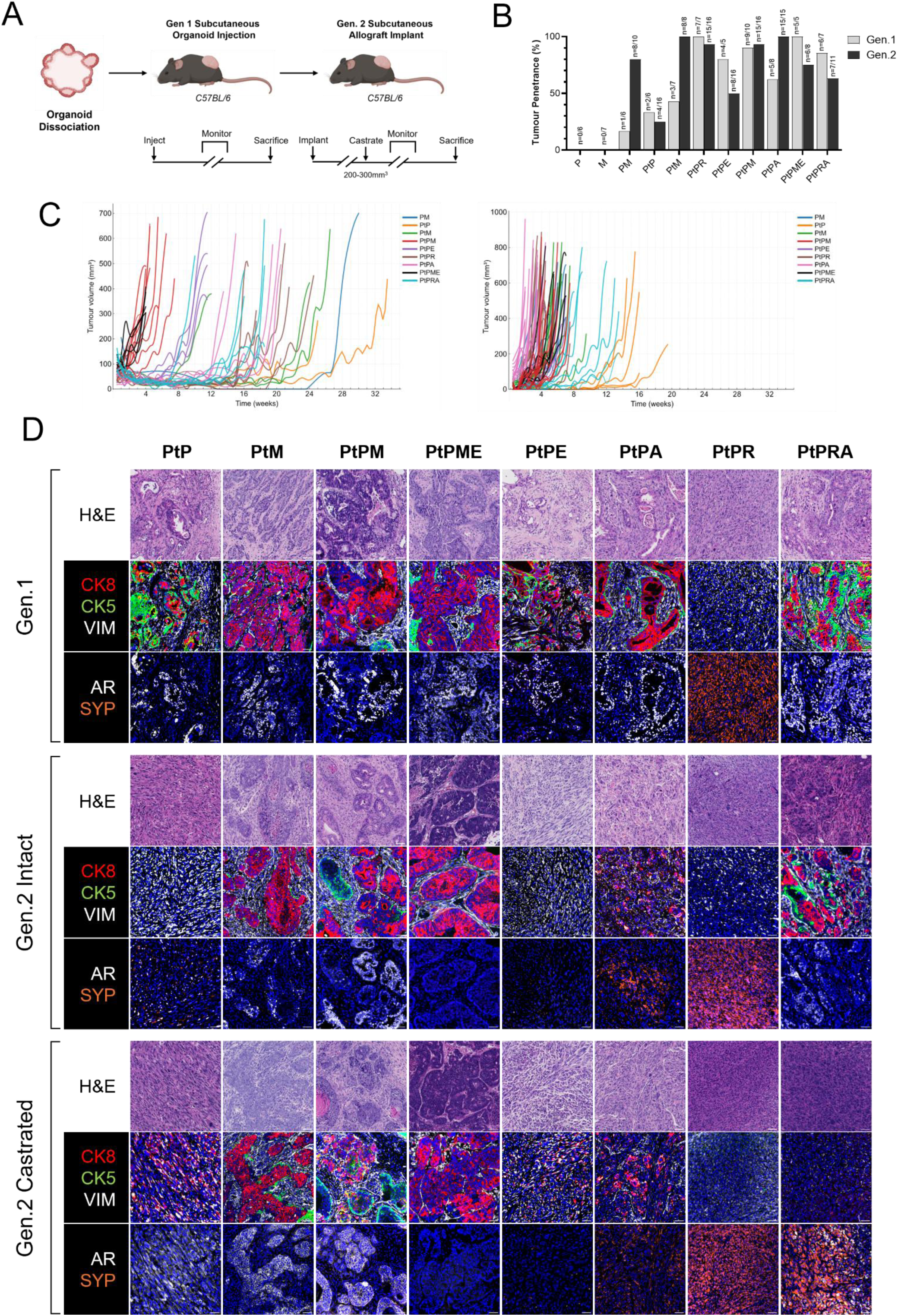
ProMPt tumours model frequent and rare CRPC subtypes. (A) Schematic representation of the strategy used to establish *in vivo* ProMPt tumour models for characterization. (B) Engraftment efficiency by genotype and generation. Cohort sizes are indicated in the figure. (C) Individual tumour growth curves per mouse, coloured by genotype, for subcutaneous Generation 1 (left) and allograft Generation 2 (right) ProMPt models. (D) Representative H&E and cyclic immunofluorescence (CyCIF) images of Generation 1 and Generation 2 tumours stained for lineage-specific markers. Serial sections were taken between H&E and CyCIF. Scale bar, 50 μm. Selected regions of interest (ROIs) and additional marker panels are shown in Figures S4 and S5.

Histopathological analysis revealed a wide spectrum of morphologies (Fig. 3D; Supp. Table 1; Supp. Figs. 4–5). The PtP model, which serves as the genetic backbone for other ProMPt combinations, resembled previously reported prostate-specific PtP GEMMs in which lineage mixed adenocarcinomas progressed toward basal and sarcomatoid carcinomas through epithelial–mesenchymal transition(36). Gen 1 PtP tumours exhibited a Ck5⁺ basal–squamous phenotype with limited Ck8⁺; Ar⁺ luminal regions, showing relatively low proliferative activity by Ki67 staining while maintaining Trop2 expression and variable Cd104 levels. As allografts, PtP tumours appeared highly cellular and sarcomatoid, expressing Cd44, consistent with transitional states indicative of basal–mesenchymal reprogramming (Supp. Fig. 4B).

**Figure 4.**
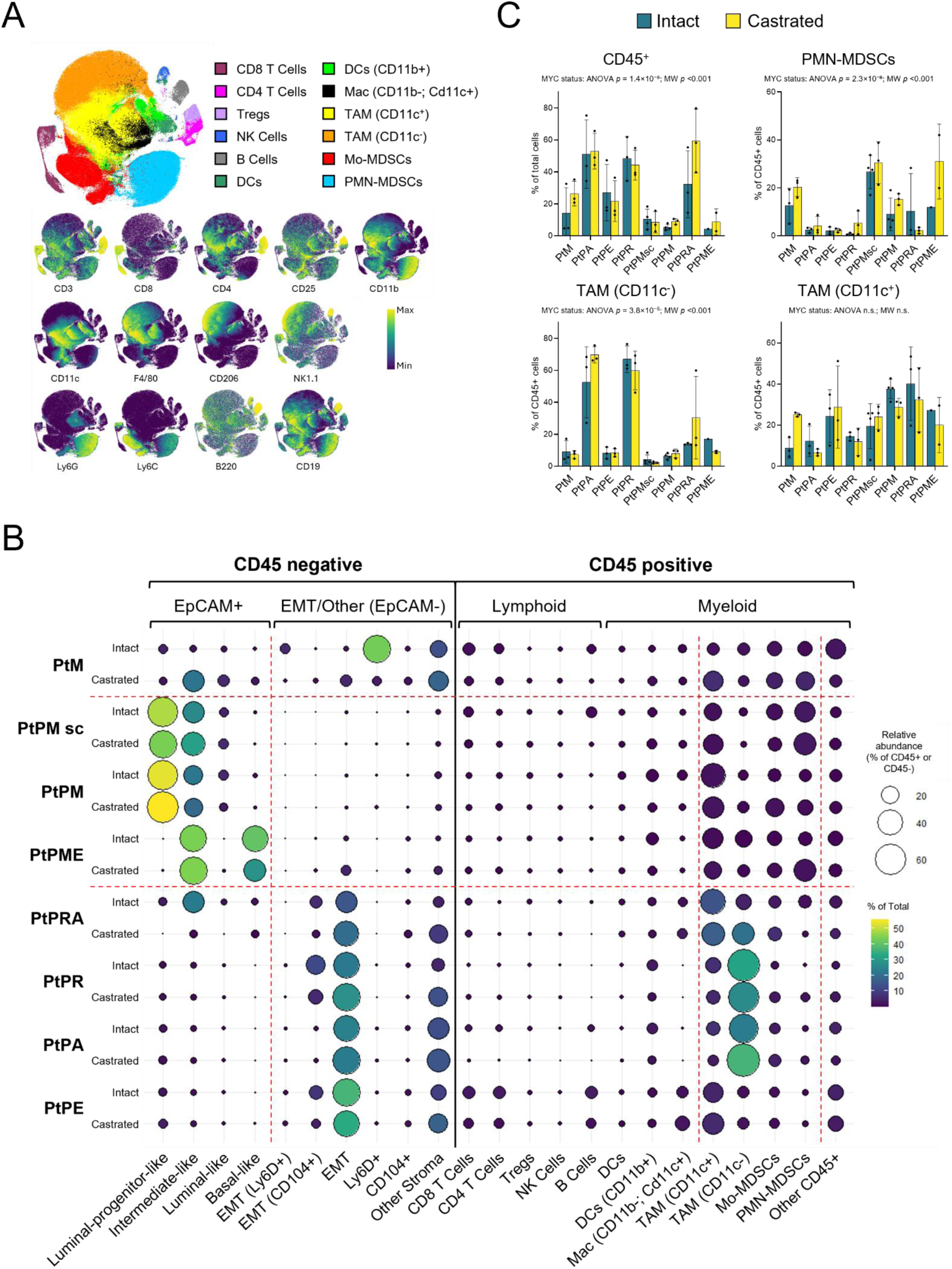
ProMPt tumours demonstrate genotype-specific immune composition. (A) t-SNE map generated from total Cd45⁺ immune cells collected across all individual tumours (n = 48), colour coded by population gate (top) or by surface marker expression detected through CyTOF. See Figure S6 for gating strategy. (B) Bubble plot showing relative abundance of cell populations segmented into Cd45⁻ and Cd45⁺ compartments for ProMPt Gen. 2 (n = 3 intact; n = 3 castrated) and PtPM Gen. 1 (n = 4 intact; n = 3 castrated) tumours. Bubble size indicates relative abundance within the Cd45⁺ or Cd45⁻ compartments, and bubble colour denotes percentage of the total population. Mean values were used for replicates; see Figures 4C and S7 for statistical analysis. Red lines delineate major phenoscapes corresponding to epithelial and immune populations. C) Bar plot showing replicate values for the indicated populations from CyTOF analysis. Two-way ANOVA was used to assess main and interaction effects of *MYC* status and castration. When overall effects were significant, pairwise Mann–Whitney U tests were performed to compare *MYC*⁺ and *MYC*⁻ groups within each condition. Bars represent mean ± SD (n = 3 per group). See also Figures S7A and S7B for extended results.

**Figure 5.**
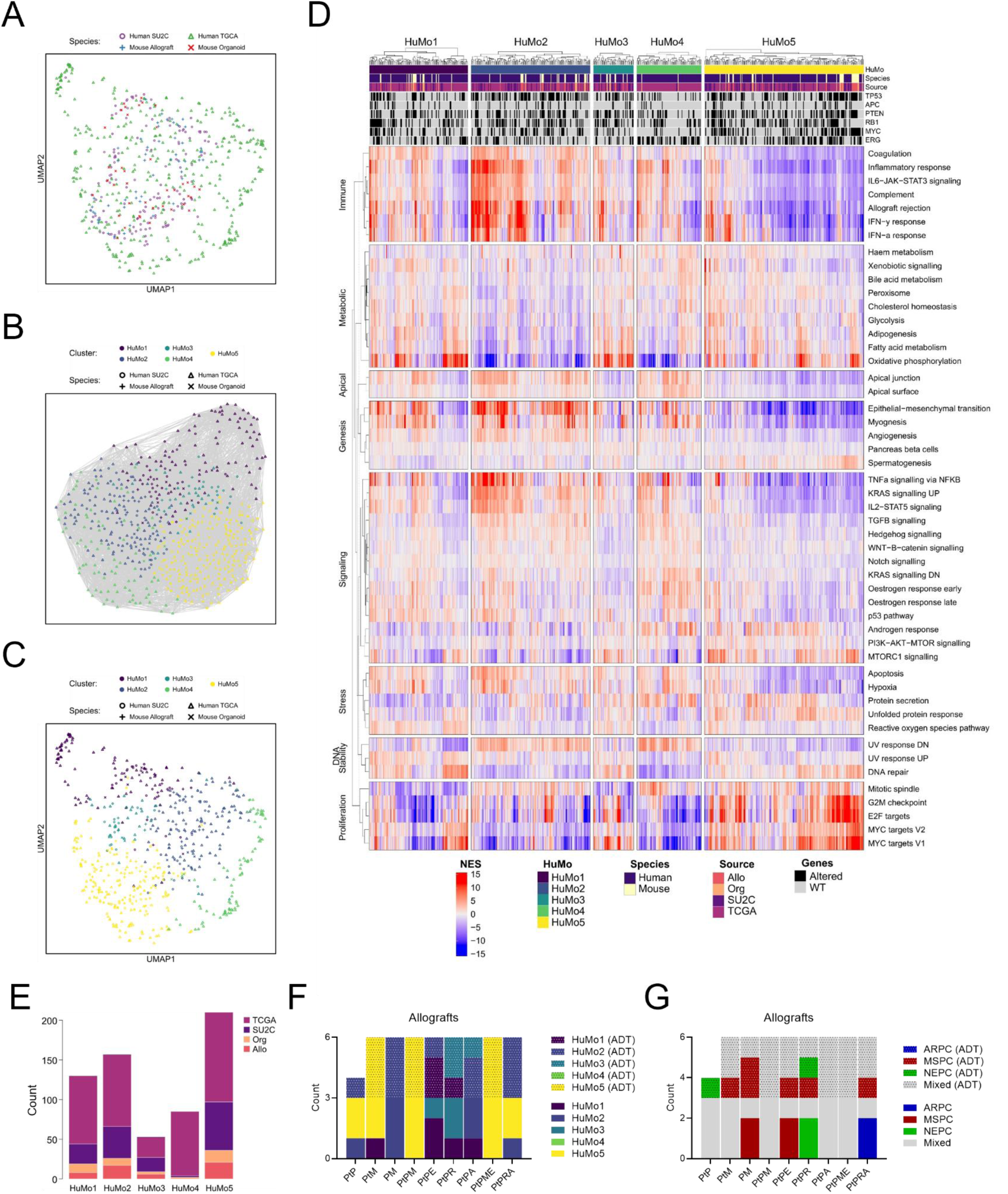
Pathway and subtype analysis of integrated human and mouse data highlights cluster specific characteristics. (A) UMAP visualisation showing intermixing of all ProMPt samples (organoids +DHT and long-term – DHT; Gen. 2 intact and castrated; n = 92) with human prostate cancer transcriptional profiles from TCGA and SU2C (n = 543). (B) Louvain clustering of the integrated ProMPt and human prostate cancer transcriptomes resolved five distinct groups. (C) UMAP visualisation of the five clusters identified, highlighting shared human–mouse (HuMo1–HuMo5) groups. (D) Hallmark pathway enrichment analysis across HuMo clusters, grouped by biological process. The scale bar indicates normalised enrichment. Species, dataset source, and genetic profiles corresponding to ProMPt-modelled aberrations are shown at the top. (E) Distribution of samples coloured by source dataset (Org: ProMPt organoids; Allo: Gen. 2 ProMPt models) within individual HuMo clusters. See also Figure S8A. (F) ProMPt allograft samples classified by HuMo cluster, coloured by cluster identity. Patterned colours indicate castrated samples (ADT). (G) ProMPt allograft samples classified into ARPC, MSPC, NEPC, and Mixed subtypes using gene signatures from Han et al.(11). Patterned colours indicate castrated (ADT-treated) samples.

In contrast, MYC-driven tumours (PM, PtM, PtPM, PtPME) consistently formed poorly differentiated adenocarcinomas. These lesions displayed glandular architectures with a well-defined Ck8⁺ luminal layer, variable Ck5⁺ basal cells, and stromal vimentin surrounding the glands (Fig. 3D). Trop2 was readily detected in luminal regions, whereas Cd104 marked basal elements, and notably, Trop2⁺Cd104⁺ double-positive cells were frequently present within glandular areas, indicative of intermediate epithelial states. The tumours were highly proliferative, retained variable Ar expression, and exhibited comparable morphology between Gen 1 and Gen 2. Some replicates, particularly in Gen 2, contained regions of squamous or sarcomatoid differentiation, and PtM tumours uniquely showed focal osseous metaplasia, features consistent with aggressive adenocarcinoma (Supp. Fig. 4A-B).

PtPE tumours exhibited progressive mesenchymal-like features from Gen 1 to Gen 2, evolving from lesions with sparse Ck8⁺; Ar⁺ luminal regions to predominantly spindle-cell, sarcomatoid morphologies. These tumours showed intense vimentin expression together with Cd104 and Cd44 positivity (Fig. 3D; Supp. Fig. 4B), consistent with EMT-like prostate cancer. The PtPR line showed a marked divergence from adenocarcinoma, characterized by sarcomatoid morphology with loss of A r, cytokeratin markers, and Trop2, together with prominent Vim and Cd44 staining. Within these tumours, interspersed regions of neuroendocrine differentiation were evident, marked by intense synaptophysin (Syp) positivity (Fig. 3D; Supp. Fig. 5). This hybrid architecture, defined by coexisting EMT- and stem-like programs, recapitulates the plastic states observed in PtPR organoids *in vitro* and mirrors their *in vivo* trajectory, in which combined *Pten*, *Trp53*, and *Rb1* loss drives transition to neuroendocrine phenotype(37,38) . These features parallel the clinical association of this genetic profile with NEPC in patients(4). The PtPA and PtPRA models produced the most heterogeneous tumours across generations, encompassing areas of poorly differentiated adenocarcinoma with Trop2⁺ luminal and Ck8⁺ glandular regions, which became less prominent in Gen 2 and following castration. These coexisted with zones of squamous differentiation, sarcomatoid morphology, and focal Syp-positive neuroendocrine areas (Fig. 3D; Supp. Fig. 5). This broad spectrum of differentiation reflects pronounced lineage plasticity and the coexistence of EMT- and stem-like programs.

Together, these results demonstrate that ProMPt tumours reproduce both prevalent prostate cancer subtypes, including prostate adenocarcinoma (PRAD), MSPC and NEPC, as well as rarer variants such as squamous and sarcomatoid carcinoma. Tumour architecture and lineage composition were primarily dictated by genotype, with *MYC* overexpression in *Pten*-null backgrounds consistently yielding aggressive adenocarcinomas, whereas combined loss of *Pten*, *Trp53*, and *Rb1* reaffirmed the previously reported predisposition to neuroendocrine differentiation. This genotype-driven heterogeneity, revealed through *in vivo* adaptation, highlights the utility of ProMPt as a platform for modelling the full spectrum of prostate cancer phenotypes in an immune competent setting.

### ProMPt tumours recapitulate CRPC immune landscapes

To complete the ProMPt characterisation pipeline, we profiled the cellular composition of ProMPt tumours at single-cell resolution by CyTOF. Pooled allograft samples from all genotypes, collected at endpoint under intact or castrated conditions, were analysed using a 23-antibody panel combined with mass-tag barcoding (Fig. 4A). t-SNE visualisation of the Cd45⁺ compartment revealed discrete clusters corresponding to major immune populations, with gating strategy performed as shown in Supplementary Fig. 6. Within the myeloid compartment, differential expression of Cd11b, Cd11c, and F4/80 delineated five principal populations, in line with previous reports in prostate and other solid tumours(6,39–44): canonical tumour-associated macrophages (TAM; Cd11c⁻), TAM (Cd11c⁺), classical dendritic cells (DCs), Cd11b⁺ dendritic cells (Cd11b⁺ DCs), and macrophages (Mac; Cd11b⁻ Cd11c⁺).

Global mapping of immune profiles revealed two major, distinct tumour microenvironmental states largely defined by *MYC* status (Fig. 4B). Tumours harbouring the *MYC*-overexpressing allele displayed an immune-restricted microenvironment characterised by low Cd45⁺ infiltration, whereas non-*MYC* tumours were immune-permissive and heavily infiltrated (Fig. 4C). Analysis of PtPM Gen 1 and Gen 2 tumours revealed comparable immune compositions, indicating stable preservation of the microenvironment upon serial passaging. Across ProMPt models, myeloid populations represented the dominant immune compartment and displayed the most pronounced genotype-specific variation. In *MYC⁺* tumours, both polymorphonuclear (PMN-) and monocytic (Mo-) myeloid-derived suppressor cells (MDSCs) were relatively more abundant than in *MYC*⁻ models (Supp. Fig. 7A). These tumours also contained a higher proportion of TAM (Cd11c⁺) relative to canonical TAM (Cd11c⁻). These findings point to an expansion of immunosuppressive myeloid populations and the establishment of an immune-restrictive microenvironment driven by *MYC* activation(45,46), in line with previous observations in prostate adenocarcinoma(47,48). Conversely, *MYC*⁻ tumours were characterised by dense macrophage infiltration, with variable TAM (Cd11c⁻)/TAM (Cd11c⁺) ratios across genotypes. Canonical TAMs (Cd11c⁻) were particularly enriched in specific contexts such as PtPA and PtPR, consistent with evidence linking mesenchymal and EMT-like states to macrophage recruitment(49). Other immune compartments exhibited more modest differences, and castration had no discernible effect across models at endpoint.

We next profiled the Cd45⁻ compartment (Fig. 4B, right; Supp. Fig. 7B). Gating on EpCAM, Trop2, Ly6D, Cd104, and Cd44 resolved epithelial and mesenchymal populations. Within the EpCAM⁺ fraction, Trop2⁺ luminal-like cells included subsets co-expressing the progenitor marker Ly6D (luminal-progenitor-like)(50), or the basal marker Cd104 (intermediate-like), while EpCAM⁺Cd104⁺ cells defined a basal-like population. In contrast, EpCAM⁻ cells were largely Cd44⁺, consistent with an EMT-like identity. The analysis was in close agreement with our spatial data (Fig. 3D; Supp. Figs. 4–5), showing clear luminal enrichment in *MYC*⁺ tumours and, conversely, pronounced EMT features in non-*MYC* allografts (Fig. 4B; Supp. Fig. 7B). Notably, PtPM models displayed a predominance of luminal-progenitor-like and intermediate-like cells, whereas PtPME tumours were enriched for intermediate- and basal-like populations.

Collectively, our characterization demonstrates that ProMPt CRPC models converge on two distinct immune–epithelial configurations. *MYC* positive tumours, which mirror classical PRAD, exhibit an immune-restricted state with expansion of regulatory and suppressive myeloid subsets. In contrast, *MYC*^-^ tumours, resembling EMT-enriched variants, display an immune-permissive, mesenchymal-shifted organisation dominated by inflammatory macrophages. The delineation of these archetypes provides a basis for linking epithelial lineage and immune contexture, enabling the identification of therapeutic vulnerabilities and supporting future immunotherapy testing.

### Cross-species transcriptomic mapping reveals conserved prostate cancer archetypes

To assess the translational relevance of the ProMPt platform and its alignment with the molecular diversity of human prostate cancer, we performed cross-species transcriptomic integration of ProMPt organoids and allografts with human datasets from TCGA (primary) and SU2C (metastatic CRPC) using a transcriptional alignment and clustering framework recently developed for hepatocellular carcinoma (HCC)(18). Uniform manifold approximation and projection (UMAP) combined with Louvain community detection identified five reproducible human–mouse (HuMo) clusters (HuMo1–HuMo5) encompassing both species and capturing conserved molecular programmes of prostate cancer (Fig. 5A–C). The joint manifold showed extensive intermixing of murine and human samples, indicating that ProMPt models occupy transcriptionally relevant disease states rather than forming mouse-specific branches. Similar to observations in HCC, mutational status alone did not fully predict transcriptional proximity or pathway activation, underscoring the functional convergence of distinct genotypes within shared signalling networks. Pathway-level analysis further resolved these five clusters according to gradients of proliferation (MYC targets, E2F, G2M), androgen response, EMT/TGFβ, and hypoxia/angiogenesis, with immune/inflammatory, PI3K–AKT, and MAPK/KRAS pathways also showing consistent cross-species alignment (Fig. 5D). HuMo1–3 contained a balanced mixture of TCGA and SU2C tumours, HuMo4 consisted predominantly of TCGA primary cancers, and HuMo5 was enriched for SU2C metastatic CRPC samples (Fig. 5E; Supp. Fig. 8A). At one end of this spectrum, HuMo5 encompassed tumours spanning the AR-active to MYC-driven continuum and represented the largest cluster overall. It exhibited the highest MYC, E2F, and G2M signalling activity, together with elevated mTORC1 and PI3K–AKT pathways, while EMT/TGFβ, KRAS, and inflammatory programmes were largely suppressed. This transcriptional profile is consistent with an androgen receptor signalling inhibitor (ARSI)-focused therapeutic strategy, potentially combined with approaches that exploit MYC-associated synthetic vulnerabilities or enhance immune engagement within the tumour microenvironment. Subtype classification, based on Han *et al.*(11), was enriched for ARPC and Mixed signatures (Supp. Fig. 8C). PtM, PtPM, and PtPME models co-localised within this cluster (Fig. 5F), consistent with their MYC-programs upregulation, PRAD histology and immune-restricted phenotype observed by CyCIF and CyTOF respectively.

At the opposite end, HuMo1 and HuMo2 defined inflamed, mesenchymal, and AR-attenuated states distinguished by divergent DNA-damage and metabolic pathway activity. Both clusters exhibited high EMT/TGFβ, hypoxia/angiogenesis, and MAPK/TNF/STAT signatures, whereas androgen-responsive, MYC/E2F, and mTORC1 programmes were comparatively low. Subtype classification across these clusters showed the greatest enrichment for MSPC tumours, with HuMo2 containing the highest proportion of WNT/STEM-like profiles(9) (Supp. Fig. 8C,E). This transcriptional space corresponded closely to the CyTOF-defined immune-rich, myeloid-heavy microenvironment, with most PtPE, PtPR, PtPA, and PtPRA allografts, displaying mixed or sarcomatoid histology, mapping within these groups (Fig. 5F).

HuMo4 was composed almost exclusively of TCGA samples, with few SU2C and no allografts, indicating limited representation of advanced disease (Fig. 5E). It reflected a primary-like cluster characterised by elevated AR activity, moderate proliferation, and mixed stem, EMT, and metabolic signatures, consistent with an intermediate disease state. HuMo3 bridged the transition between these primary-like and advanced states, combining high proliferative (E2F/MYC) activity with moderate EMT and variable immune activation. This cluster contained a higher proportion of SU2C samples relative to HuMo4. Notably, several PtPR allografts, displaying immunohistochemical evidence of neuroendocrine differentiation, mapped to HuMo3 and were classified as NEPC by subtype analysis (Fig. 5G), consistent with their CyCIF profiles.

In summary, cross-species HuMo integration identified two dominant CRPC transcriptional archetypes that occur in both human and mouse prostate cancer. One archetype (HuMo5) exhibited AR and MYC activity, epithelial phenotype and low immune infiltration. The second (HuMo2) was characterised by EMT, inflammatory signalling and a macrophage-rich microenvironment. ProMPt tumours distributed along this axis: luminal *MYC*⁺ allografts aligned with the immune-restricted HuMo5 state, whereas *MYC*^-^ sarcomatoid tumours with prominent macrophage infiltration mapped to HuMo2. By modelling both endpoints within a genetically controlled, immunocompetent system, ProMPt enables mechanistic dissection of epithelial–immune interactions and rational testing of therapies tailored to genotype and lineage state.

### Media permutation exposes genotype-specific androgen–EGF dependency

Building on the HuMo and *in vivo* analyses that pointed to distinct signalling dependencies across genotypes, we next sought to test how microenvironmental cues influence response to androgen deprivation and pathway activation. Rather than performing a traditional compound screen, we implemented a media-permutation strategy to interrogate the requirement for ligand-driven signalling in organoid adaptation (Fig. 6A). Seven ProMPt lines were cultured in triplicate with or without DHT, and within each condition, individual components of standard prostate organoid media, EGF, A83-01, Noggin, and R-Spondin, were withdrawn. In total, 210 organoid cultures were multiplexed using TOB*is* mass-tag barcoding and profiled by CyTOF with a 34-antibody panel capturing epithelial states, cell-cycle dynamics, and key signalling nodes(19) .

**Figure 6.**
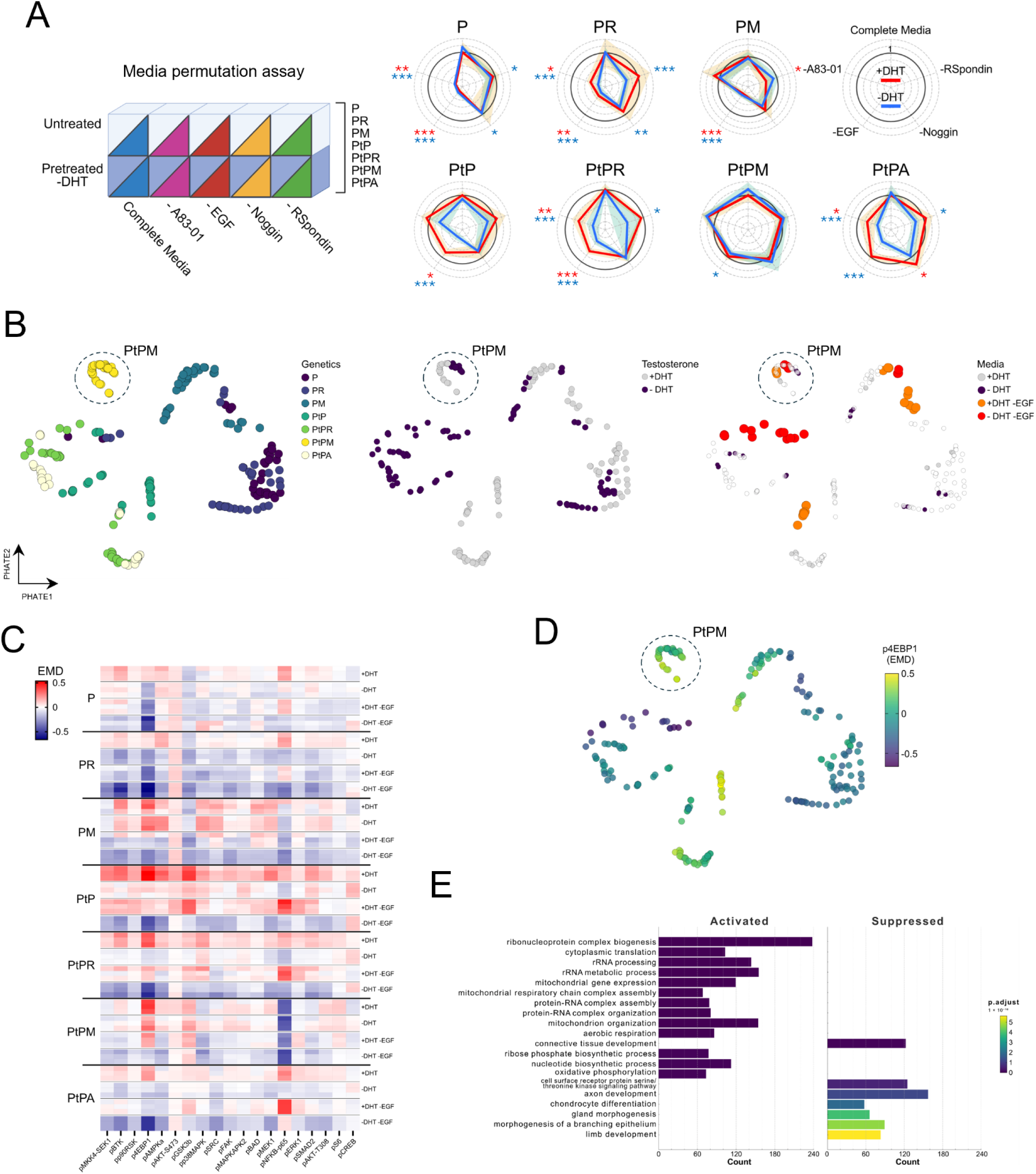
Media permutation exposes genotype-specific androgen–EGF dependency. (A) Left: Schematic of the media permutation strategy in ProMPt organoids. The indicated ligands were individually removed from complete media under untreated (+DHT) or long-term testosterone-deprived (–DHT) conditions. Right: Radar plots showing relative viability for each line and ligand-removal condition, with culture status indicated in the top-right plot (red, +DHT; blue, –DHT). Shaded regions (orange, light blue) represent standard deviation across six replicates. Statistical significance determined by Welch’s t-test comparing each ligand-removal condition with its matched control (\**P* < 0.05, ** *P* < 0.01, *** *P* < 0.001). (B) Earth mover’s distance (EMD)–PHATE embedding of organoid cultures from the media permutation assay, coloured by genotype (left), DHT status (middle), and ligand permutation condition (right). Each dot represents the EMD–PHATE position. Three biological replicates per condition. (C) EMD heatmap of phospho-protein profiles across genotypes and culture conditions (DHT and EGF deprivation). Biological triplicates are shown. (D) EMD-PHATE coloured according to p4E-BP1 EMD score. (E) Top 20 enriched GO_BP pathways from bulk RNA-seq comparison of *MYC⁺* versus *MYC⁻* ProMPt organoids. Analysis includes 21 models (*MYC⁻* = 13; *MYC⁺* = 8). P values were adjusted for multiple testing.

**Figure 7.**
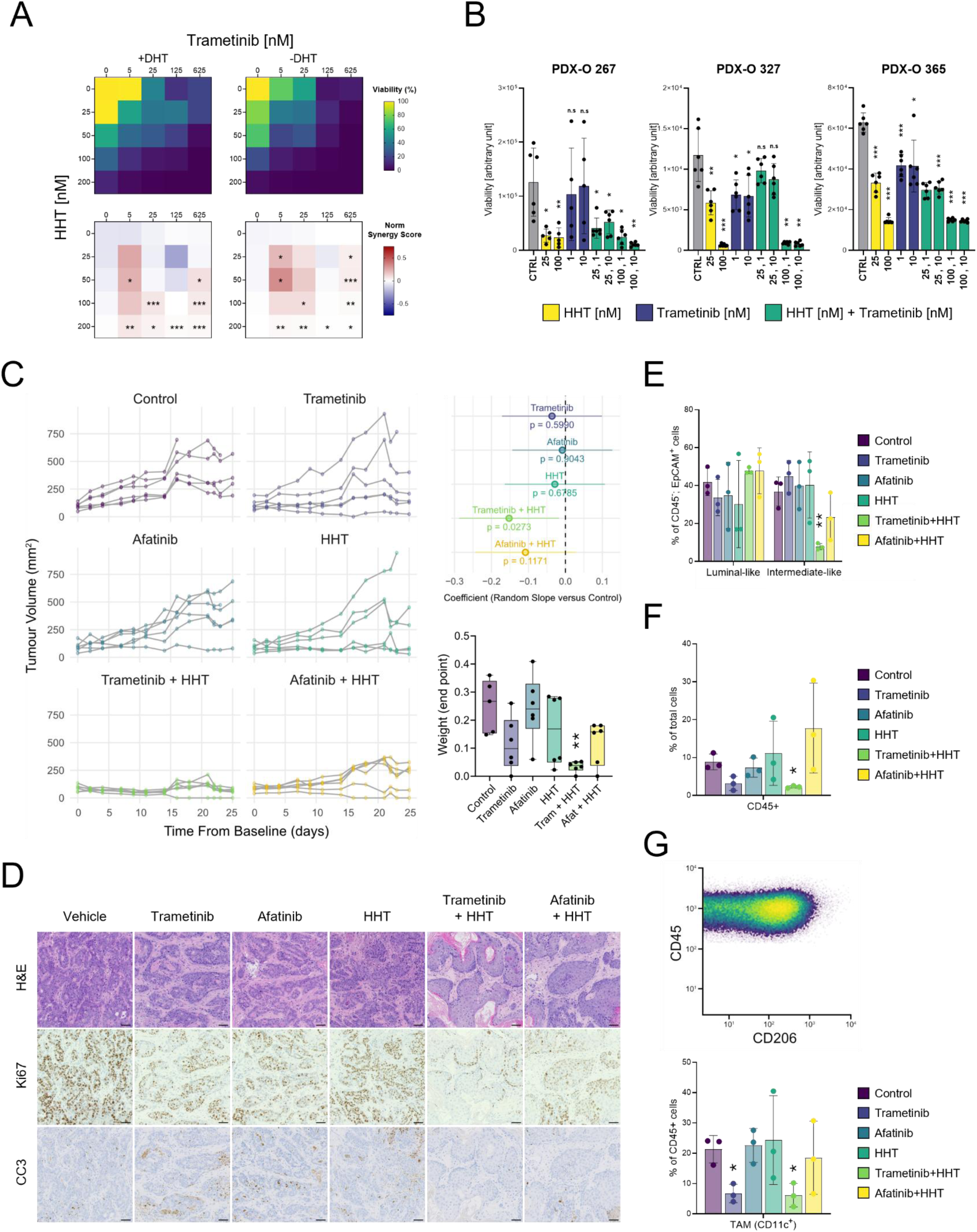
Inhibition of MAPK and Protein Translation synergize to inhibit tumour development. (B) Viability heatmap (top) and Bliss synergy score map (bottom) for PtPM organoids treated under +DHT or –DHT conditions with trametinib in combination with homoharringtonine (HHT). A two-sided one-sample t-test was applied to triplicate inhibition fractions at each dose pair to assess deviation from Bliss additivity. Asterisks denote dose pairs where observed inhibition significantly exceeded the Bliss expectation (\**P* < 0.05, \*\**P* < 0.01, \*\*\**P* < 0.001). (C) Relative viability of PDX-derived organoids following treatment with homoharringtonine (HHT), trametinib, or the combination. Data represent six replicates per condition. Welch’s t -test was used to compare each treatment with the control (\**P* < 0.05, \*\**P* < 0.01, \*\*\**P* < 0.001). Bars indicate mean ± SD. (D) Tumour growth curves (left) for PtPM subcutaneous tumours generated by organoid injection across six treatment arms. Each line represents one mouse (n = 6 per group). A mixed-effects regression model was used to assess differences in growth between treatment and control cohorts (top right). Final tumour weights (bottom right) were compared with vehicle controls using Welch’s t-test (\*\**P* < 0.01). Treatments were administered over a 25-day time course. (E) Representative histology and immunohistochemistry staining of PtPM tumours collected at endpoint (n = 3). Scale bar, 50 μm. (F) Relative abundance of luminal-like and intermediate-like cells in endpoint PtPM tumours across treatment arms, quantified by CyTOF analysis. Comparisons with vehicle controls were performed using Welch’s t-test (\*\**P* < 0.01). Bars indicate mean ± SD (n = 3 per group). (G) Abundance of CD45^+^ in endpoint PtPM tumours across treatment arms, quantified by CyTOF analysis. Comparisons with vehicle controls were performed using Welch’s t-test (\**P* < 0.05). Bars indicate mean ± SD (n = 3 per group). (H) Density plot showing CD206 expression in TAMs (CD11c⁺) from pooled CyTOF data of the PtPM treatment experiment (top). Bottom: relative abundance of TAMs (CD11c⁺) in endpoint PtPM tumours across treatment arms. Comparisons with vehicle controls were performed using Welch’s t-test (\**P* < 0.05). Bars indicate mean ± SD (n = 3 per group).

Viability profiling across all conditions (Fig. 6A) revealed that EGF withdrawal exerted the strongest effect, markedly reducing proliferation across multiple genotypes, particularly those with intact *Pten*. In PtP, PtPR, and PtPA organoids, removal of EGF was partly tolerated under DHT-supplemented conditions but became strongly detrimental upon testosterone withdrawal, unmasking a latent hormone requirement concealed under complete media. In contrast, PtPM organoids maintained viability across all conditions, indicating that MYC activation confers resistance to growth-factor deprivation, specifically to EGF loss and associated MAPK pathway inhibition. To assess this relationship directly, we pharmacologically tested four *MYC*-expressing and *MYC*-deficient organoid pairs using the MEK inhibitor trametinib. *MYC^+^* organoids displayed greater resistance to trametinib than their *MYC* negative counterparts (Supp. Fig. 9A), establishing MYC as a determinant of MAPK - inhibition tolerance.

Integration of Earth Mover’s Distance (EMD) analysis of all CyTOF profiles using PHATE embedding (Fig. 6B) showed that samples segregated primarily by genotype and DHT status, with EGF withdrawal producing the most pronounced global shift across ligand-permutation conditions. Most genotypes underwent marked reorganisation of their cell-state composition upon EGF removal, whereas PtPM remained tightly clustered across all conditions, consistent with its stable viability. Lineage analysis revealed, across genotypes, two principal epithelial populations: a luminal -like Trop2⁺Cd104⁻ fraction and a Trop2^mid^Cd104⁺ fraction consistent with an intermediate-like profile (Supp. Fig. 9B). Two-way ANOVA demonstrated significant effects of genotype and treatment on the luminal-like population, which decreased sharply upon DHT withdrawal but increased significantly in some genotypes following EGF removal (Supp. Fig. 9C). Intermediate-like cells showed the opposite trend, expanding modestly under androgen deprivation but becoming markedly diminished upon EGF withdrawal, a pattern largely conserved across models but attenuated in PtPM. The balance between these two populations indicates a reciprocal regulatory relationship, whereby androgens maintain luminal identity while EGF signalling stabilises intermediate states.

To better understand the mechanisms underlying PtPM’s resistance to growth-factor and hormone withdrawal, we next examined the phospho-signalling data. As expected, phospho-AKT levels mirrored *Pten* status, whereas phosphor-BTK, phospho-4EBP1 (p4EBP1), phospho-GSK3β, pMEK, and phospho-P65 exhibited the greatest variability across genotypes and conditions (Fig. 6C; Supp. Fig. 9D). Among these, p4EBP1 was particularly notable as its levels decreased sharply under combined DHT and EGF withdrawal in most models but remained largely sustained in PtPM. Visualisation of p4EBP1 across the PHATE embedding (Fig. 6D) and confirmatory gating analyses (Supp. Fig. 9E–F) revealed that p4EBP1 expression was only minimally reduced in PtPM across experimental conditions, with enrichment in intermediate-like populations but maintained expression in both epithelial compartments. These findings indicate that *MYC*, in a *Pten*-deficient background, sustains translational activity under growth-factor stress, providing a mechanistic basis for PtPM’s resistance to EGF deprivation and MAPK inhibition. Given the well-established role of *MYC* in transcriptional regulation of the protein synthesis machinery(51,52), we next examined whether this link was reflected at the transcriptomic level across ProMPt organoids. Consistent with this model, bulk RNA-sequencing comparing *MYC*-positive and *MYC*-deficient ProMPt organoids revealed top enrichment for translation-related pathways (Fig. 6E), and analysis of SU2C and RMH patient cohorts confirmed a strong correlation between MYC-targets and Reactome Translation signatures (Supp. Fig. 9G).

In summary, the multidimensional TOBis mass cytometry–based media permutation analysis in ProMPt organoids enabled systematic dissection of how androgen and growth-factor cues shape epithelial responses and uncover context-specific dependencies masked by standard culture conditions. EGF withdrawal, modelling MAPK inhibition, acted synergistically with androgen deprivation, yet *MYC* overexpression sustained viability under dual stress in a *Pten*-deficient background. Sustained 4E-BP1 phosphorylation identified protein translation as a potential mechanism of resistance. Together, these findings show that androgens and EGF cooperatively preserve epithelial viability in organoid settings, whereas combined *Pten* loss and *MYC* activation maintain translational output under stress, reinforcing translational control as a MYC-linked vulnerability within defined genetic contexts.

### Co-targeting MAPK Signalling and Protein Translation Reveals a Therapeutic Vulnerability

Recent studies have demonstrated the therapeutic activity of MAPK inhibition in prostate cancer models *in vivo*(53), and MEK inhibition with trametinib has progressed to Phase II trials in castration-resistant disease (NCT01990196, NCT02881242). However, our HuMo analysis and media-permutation CyTOF experiments indicated that PtPM ProMPt models maintain translational output and tolerate MAPK inhibition, suggesting that concurrent targeting of translation may overcome this resistance. We therefore focused on the translational machinery and selected homoharringtonine (HHT), an FDA-approved inhibitor of protein synthesis, as a clinically tractable option(54).

To identify rational combinations, we performed a preliminary screen testing HHT with trametinib, afatinib, or capivasertib, selected to probe the PI3K–AKT-mTOR signalling axis, across PtP, PtM, PM and PtPM organoids (Supp. Fig. 10A). Interestingly, HHT combined with either trametinib or afatinib showed a trend toward synergistic effects across several ProMPt lines, whereas HHT with capivasertib offered no benefit over monotherapy. Among genotypes, PtPM organoids were comparatively less sensitive to trametinib, but exhibited a greater response when trametinib was combined with HHT, in line with their elevated p4EBP1 levels and high translational capacity. Based on this data, we prioritised PtPM for further analysis and evaluated the trametinib + HHT combination under both +DHT and −DHT conditions (Fig. 7A). In both settings, the combination reduced growth and produced moderate synergy, indicating that the effect is largely independent of androgen signalling.

To explore translational relevance, we next tested the combination in three CRPC organoid lines generated from PDX (PDX-O)(55), representing distinct genetic backgrounds. The most proliferative line, PDX-O 267, which carries *MYC* amplification with *PTEN* and *TP53* loss (Supp. Table 4), was highly sensitive to HHT and showed the greatest growth suppression (Fig. 7B; Supp. Fig. 10B). In contrast, trametinib alone had only minor effects across all three lines, and no clear synergism with HHT was observed in these human models, likely reflecting the greater genetic complexity and distinct signalling dependencies inherent to these patient-derived CRPC organoids.

We next validated the treatment *in vivo* using the PtPM model. All mice were subjected to androgen deprivation (degarelix) and treated with trametinib, afatinib, or HHT as single agents, or with the combinations trametinib + HHT or afatinib + HHT for four weeks (Supp. Fig. 10C). Single-agent regimens produced limited responses, whereas trametinib + HHT induced marked tumour regression and afatinib + HHT showed a milder but consistent trend (Fig. 7C; Supp. Fig. 10D). Histological analysis at end point revealed extensive epithelial remodelling in both combination groups, with tumours showing a squamous-like morphology and a marked reduction in proliferation with minimal apoptosis at this late time point (Fig. 7D).

To further investigate how the treatment affects both the tumour and its microenvironment, we profiled endpoint tumours by single-cell CyTOF using the same 34-marker panel described in Fig. 4. This analysis confirmed a pronounced reduction in epithelial content, which was most significant in the trametinib + HHT cohort (Supp. Fig. 11A). In PtPM tumours, epithelial cells were uniformly Trop2⁺ and segregated into Ly6D⁺ luminal-progenitor-like and Cd104⁺ intermediate-like states (Supp. Fig. 11B). The combination preferentially depleted the intermediate-like population, the same compartment that exhibited elevated p4EBP1 in organoid assays (Fig. 7E). Analysis of the immune compartment revealed a marked decrease in Cd45⁺ infiltrates following trametinib treatment, both as single agent and in combination with HHT (Fig. 7F). Although T-cell and MDSC abundance remained largely unchanged, tumour-associated macrophages (Cd11c⁺), characterised by high Cd206 expression consistent with an M2-like polarisation state, were strongly reduced in trametinib treated tumours (Fig. 7G; Supp. Fig. 11C,D). These data indicate that the effective response to trametinib + HHT involves perturbation of both tumour-intrinsic epithelial states and the surrounding microenvironment, potentially explaining the lack of synergism in PDX-O models.

Together, these data show that dual MEK and translation inhibition effectively targets the MYC-driven, androgen-independent state uncovered by the ProMPt platform. The combination exerts complementary effects: within the tumour epithelium, it selectively eliminates the translationally active intermediate population; within the microenvironment, it reduces M2-like macrophages that support tumour growth. This coordinated remodelling highlights protein-synthesis control as a tractable MYC-linked vulnerability and suggests that co-targeting translation and MAPK signalling may represent a rational therapeutic strategy for aggressive, MYC-high prostate cancer.

## DISCUSSION

Prostate cancer progression is shaped by the interplay between genetic complexity, lineage plasticity and microenvironmental signalling, yet existing experimental systems rarely enable parallel interrogation of these dimensions at scale. ProMPt was designed to overcome these limitations by utilizing a novel panel of genetically defined, multiallelic organoids that are amenable to rapid transplantation in immunocompetent hosts. This enables scalable and parallel evaluation of genotype-specific therapy responses, emergent epithelial states, and resulting tumour–immune interactions. ProMPt organoids maintain robust Ar signalling and transcriptomic profiling supported derivation of a murine Ar-response signature ideal for interpreting Ar activity across murine datasets. However, androgen deprivation alone did not induce durable lineage switching or compromise viability, consistent with evidence that plastic transitions require additional stromal and niche-derived cues(5). A media-permutation assay showed that modifying niche components can unmask otherwise latent androgen dependencies, making specific genotypes sensitive to loss of Ar signalling. These data highlight the importance of microenvironmental inputs in regulating responses to selective pressure and illustrate how controlled modulation of culture conditions can provide a more dynamic *in vitro* system for studying prostate cancer evolution. The current ProMPt organoid biobank accounts for most major genomic drivers of advanced prostate cancer but is intrinsically expandable. Because recombination occurs from endogenous floxed alleles using Ubc-CreERT2 and 4-OHT, organoids are derived without viral transduction or exogenous CRISPR editing, avoiding selection bottlenecks and preserving early-passage clonal integrity. Additional combinations can be introduced through breeding or by layering Cas9–sgRNA ribonucleoprotein editing onto defined genotypes(56), providing also a controlled framework to examine the impact of mutation sequence on tumour progression. ProMPt is compatible with targeted or genome-wide CRISPR perturbation studies and pooled barcoding approaches such as PRISM(57,58), enabling multiplexed drug sensitivity or fitness screens across genetically defined contexts. Because organoids are established from purified basal or luminal populations prior to recombination, ProMPt can also interrogate cell-of-origin effects, a critical determinant of tumour phenotype, as identical genetic lesions can have distinct effects depending on whether they arise in luminal progenitors or basal cells(56,59).

Across genotypes, ProMPt tumours developed into CRPC and recapitulated the major phenotypes of advanced human disease, including classical adenocarcinoma, mixed glandular–squamous lesions and poorly differentiated or sarcomatoid morphologies. It is important to note that, despite the breadth of genotypes modelled, a subset of ProMPt lines failed to engraft *in vivo*. As subcutaneous engraftment was chosen for scalability, it is possible orthotopic implantation, stromal co-transplantation or androgen supplementation may improve engraftment efficiency. Multidimensional profiling revealed two dominant progression patterns associated with distinct epithelial and immune states. We found that tumours arising in a *Pten*-knockout, *MYC*-overexpressing background retained glandular architecture, exhibited high proliferative activity and maintained luminal programmes. These features were associated with an immune-restricted microenvironment enriched in macrophages and myeloid-derived suppressor cells, a landscape characteristic of human CRPC, where therapeutic strategies targeting myeloid populations have shown promising activity in patients(60). In contrast, tumours in a *Pten*; *Trp53* double-knockout background lacking *MYC* frequently downregulated luminal markers and acquired EMT/sarcomatoid or MSPC features. In models such as the PtPR, they also displayed focal neuroendocrine differentiation, consistent with prior reports (6,38). These tumours supported a more immune-permissive microenvironment, including extensive tumour-associated macrophages, as observed in certain metastatic CRPC cases and in preclinical models (6,46,61,62). To what extent immune composition is dictated by oncogenic context versus histopathological subtype remains to be explored. Notably, cross-species HuMo transcriptomic mapping(18) highlighted the translational relevance of ProMPt, and a majority of our tumours and CRPC patient samples converged between two molecular archetypes that reflected similar findings from our pathology and TME characterization. Low AR and MYC activity with an enrichment of EMT and stem-associated programmes, and an inflamed immune state defined one major archetype. Comparatively the other retained variable AR activity, MYC signalling, PI3K-mTOR activation and was associated with reduced inflammatory activity. Together, results of this expansive characterization converge across assays and demonstrates the potential of the ProMPt platform to facilitate therapeutic investigation tailored to specific subtypes.

To identify therapeutic vulnerabilities, we utilized TOBis barcoding to profile organoid cultures at scale under media permutation conditions using single-cell mass cytometry(63). Non-*MYC* genotypes showed a marked dependency on EGF, reinforcing MAPK signalling as a therapeutic target(64,65) and consistent with the evaluation of MEK inhibition in prostate cancer (NCT01990196, NCT02881242). In contrast, PtPM organoids lacked this dependency, suggesting MYC contributes to resistance to MAPK pathway inhibition, as observed in other solid tumours (66,67). Integration of phospho-proteomic and transcriptional data indicated that PtPM organoids preserve protein translation, despite androgen and EGF withdrawal. This is consistent with a molecular programme in which *Pten* loss sustaining PI3K–AKT–mTOR activity and phosphorylation of 4EBP1(68), *MYC* amplification boosts ribosome biogenesis and translational output(51), and diminished Ar signalling removes translational restraint(69). Together, these alterations can maintain protein synthesis despite upstream pathway inhibition, offering a plausible rationale for tolerance to MEK inhibition. We also observed epithelial state–specific effects: luminal-like cells were sensitive to androgen withdrawal, whereas intermediate cells were preferentially sensitive to EGF withdrawal. In PtPM organoids, intermediate cells retained higher levels of phosphorylated 4EBP1, consistent with cell type specific translational control(70). Our current epithelial state annotation is based on a focused set of markers (Trop2, Ly6D and Cd104), which will refine as additional single-cell atlases and functional analysis resolve epithelial hierarchies across more models. Protein translation is emerging as an attractive therapeutic target in aggressive prostate cancer(71,72). Given our findings, we tested whether dual inhibition of MAPK signalling and translation using homoharringtonine, a translation elongation inhibitor approved for the treatment of chronic myeloid leukaemia resistant to tyrosine kinase inhibitors(54), could enhance therapeutic efficacy. In the castration resistant PtPM models, the combination outperformed either monotherapy, markedly reducing tumour burden, depleting the intermediate cell compartment and decreasing M2-like macrophages. MEK inhibition has been reported to impact M2 macrophages(73,74), providing a plausible explanation for the coordinated effects on epithelial and immune components, an observation that warrants further investigation.

Collectively, this work establishes ProMPt as a flexible and scalable platform for basic discovery and preclinical testing that integrates with multimodal profiling and functionalisation strategies, and can be extended to other tumour types.

## Resource Availability

### Lead Contact

Further information and requests for resources and reagents should be directed to and will be fulfilled by the lead contact, Marco Bezzi.

### Materials Availability

ProMPt organoid and allograft models generated in this study have not yet been deposited in any repositories; however, these materials are available upon request.

### Data and Code Availability

- Raw and processed RNA-sequencing data will be deposited in an appropriate public repository prior to publication. Data supporting the findings of this study are available from the corresponding author upon reasonable request.
- This paper analyses publicly available datasets, accession numbers and links to the datasets are listed in the Key Resources Table.
- All other data associated with this study are available in supplemental figures and tables. Any additional information required to reanalyse the data reported in this paper is available from the lead contact upon request.
- This paper does not report original code.

## Acknowledgements

The authors are grateful to the ICR Confocal Microscopy, Flow Cytometry, and Breast Cancer Now Histopathology facilities for their support. Additional thanks to past and present staff members of the Animal Welfare and Ethical Review Body and biological safety unit of the ICR. This work was supported by Cancer Research UK (RCCFEL\100053 Career Establishment Award to M.B.; Convergence Science Centre PhD fellowship to E.F.), American Association for Cancer Research (AACR Clinical/Translational Cancer Research Fellowship 18-40-11-BEZZ to M.B.), Prostate Cancer UK (RIA21-EOI-020 Research Innovation Award to M.B.). Several schematic figures were prepared using BioRender.

## Author Contributions

Conceptualization and funding acquisition, M.B.; Methodology, N.P. and M.T.; Investigation, M.B., N.P., M.T., A.T., F.G., R.R., E.F., B.G. and I.F.; Analysis: M.B., N.P., J.H., M.T., G.S. and W.Y.; Software, J.H., G.S. and W.Y.; Writing – original draft: M.B., N.P. and M.T.; Writing – review & editing: M.B., N.P., M.T., A.S., J.S.d.B. and C.J.T.; Resources: A.J.N., M.C., J.G.C., C.J.T. and J.S.d.B.

## Declaration of AI-assisted writing

ChatGPT (OpenAI) was employed to support grammar correction and clarity improvements during manuscript drafting. All sections generated with assistance were critically reviewed and edited by the authors, who take full responsibility for the final content.

## MATERIALS AND METHODS

### Experimental model and subject details

#### Animal models

All GEMMs involved in the creation of ProMPT (see key resources table) were backcrossed to UBC-Cre-ERT2 mice (JAX, 007001) for one generation to produce Cre-ERT2 heterozygous progeny. Subsequent inbreeding of resulting generations was used to maintain Cre-ERT2 heterozygosity with the allele donated by either dam or sire. Loss of function alleles were bred to homozygous status.

Overexpression alleles were bred to heterozygous status. Full list of allelic combinations obtained through breeding is provided in Fig.1D. Genotyping was performed by Transnetyx (TN, USA) using real-time PCR. 10–12 weeks old C57BL/6 males (JAX, 000664) were used for all subcutaneous and preclinical experiments. For primary tumour engraftment studies, organoids were mechanically dissociated and filtered to obtain single-cell suspensions, mixed 1:1 with PBS and Matrigel (Corning, 356231) and injected subcutaneously into a single depilated and sterilised flank per animal. Organoid cultures were tested using a Mycoplasma PCR detection kit (Abcam, ab289834) following the manufacturer’s instruction prior to the start of the experiment. 1.5x10^6^ cells were injected for all genetic cohorts except for the PM group, for which 3x10^6 cells were injected (Fig. 3).

Allografts were generated by transplanting a 1×1×1 mm piece of freshly dissected tumour from the primary cohort into the flank of anaesthetised mice. When implemented, surgical castration was performed once tumours reached a size of 200–300 mm³. Appropriate pre- and post-operative analgesia was administered for all surgical procedures. For all experimental studies mice were euthanized when they exhibit >20% weight loss or met euthanasia requirements by exhibiting pain, distress or tumour burden.

All breeding and experimental protocols were approved and monitored by the Institute of Cancer Research Animal Welfare and Ethical Review Body (PP0127591, PP0936772), in compliance with the UK Home Office Animals (Scientific Procedures) Act 1986, the United Kingdom National Cancer Research Institute guidelines for the welfare of animals in cancer research(75), and the ARRIVE guidelines(76). Animals had access to sterilised food and water *ad libitum*, and facility conditions were maintained according to regulatory guidelines. The number of mice used for each study is included in the Figure Legends.

#### Murine prostate organoid generation and culture

Murine prostate organoids were established and cultured as previously described(27,33). Briefly, UBC-Cre-ERT2 positive male mice were aged 3 months to allow for complete prostate maturation. All prostate lobes were collected, minced and digested in Advanced DMEM/F-12 (Thermo Fisher, 12634010) supplemented with 5 mg/mL collagenase type II (Thermo Fisher, 17101015), 1x GlutaMAX (Thermo Fisher, 35050061), 10 mM HEPES (Thermo Fisher, 15630080), 1x Penicillin–Streptomycin (Thermo Fisher, 15140122) and 10 μM Y-27632 (Selleckchem, S1049) for 2 h at 37°C shaking. Cells were filtered (40 μm) and stained using CD24 (FITC; Thermo Fisher, 11-0241-82) and CD49f (Alexa Fluor 647; BD Biosciences, 562494) for 1 h on ice, followed by FACS sorting to separate basal and luminal populations (Fig.1C). Organoids were maintained in complete medium composed of Advanced DMEM/F-12 supplemented with 1x GlutaMAX, 10 mM HEPES, 1x Penicillin–Streptomycin, 1x B-27 supplement (Fisher Scientific, 11530536), 1.25 mM N-acetyl-L-cysteine (Thermo Fisher, A9165), 50 ng/mL EGF (Thermo Fisher, AF-100-15), 100 ng/mL Noggin (Thermo Fisher, 120-10C), R-spondin (10% conditioned medium or 500 ng/mL; Thermo Fisher, 120-38), 200 nM A83-01 (Selleckchem, S7692), and 1 nM DHT (Sigma, A8380). 10 μM Y-27632 was supplemented for 48 h after plating when cells were passaged (approximately every 5 days). Cre-mediated allelic recombination was induced by adding 100 nM 4-hydroxytamoxifen (Sigma, H7904) to the culture medium over two passages (10 days). Organoids with fewer than 12 passages were used for all experiments. To model Androgen Deprivation Therapy (ADT), DHT was removed from culture for 48–72 h (short-term DHT withdrawal) or 15–17 days (long-term DHT deprivation).

#### Organoid media permutation

Untreated and long-term DHT deprived organoids were used for all assays. For CyTOF analysis, 20,000 single cells were plated in 50 μL Matrigel in a 48-well plate, in triplicate. On the 3^rd^ day after plating, cells were starved for 6 h by removing EGF, Noggin, R-spondin, and A83-01 from the culture medium. Organoids were then cultured in permutated media for 48 h (Fig. 6A) before samples were prepared for CyTOF acquisition (see below). For imaging and viability assessment, organoids were starved as above for 6 h, dissociated, and 500 single cells were seeded in a 15ul Matrigel layer in a 96 well μ-Plate (Ibidi, 89646) in triplicate. Cells were plated and cultured for 5 days in permutated media. 10 μM Y-27632 was supplemented for 48 h after plating (Fig.6A).

#### Organoids therapeutic screening

To assess Trametinib efficacy in MYC stratified samples (Fig. S9A), untreated organoids were dissociated and plated in triplicate. Treatment was administered for 5 days at which time wells were imaged, and organoid area was assessed (see below). Untreated and long-term DHT deprived organoids were plated in triplicates for all combinatorial screenings. 500 single cells were seeded in a 15 μL Matrigel layer in a 96-well μ-Plate. 10 μM Y-27632 was supplemented for 48 h. Treatment started 48 h after plating and lasted 72 h before viability was assessed. During the preliminary screening, cells were treated with Homoharringtonine (HHT; MedChem Express, HY-14944; 0-50 nM) in combination with Trametinib (Cayman Chemical 16292), Afatinib (Selleckchem, S1011), or Capivasertib (Selleckchem, S8019) (0-2.5 μM) (Fig. S10A). During the following screen, cells were treated with HHT (0-200 nM) and Trametinib (0-625 nM) either alone or in combination (Fig.7A).

#### Patient-Derived organoids (PDX-Os)

PDX-Os were provided by Prof. Johann de Bono (Institute of Cancer Research). Organoids were generated from PDXs derived from human CRPC biopsies using established methods (33,55,77,78) . Briefly, biopsies were acquired from subjects whose tumours had progressed despite castration, AR pathway inhibitors (ARPIs), taxane therapy, and were implanted in nude mice to generate PDXs that were serially transplanted for propagation. Following cryorecovery, PDX-Os were seeded in Ultimatrix (Bio-Techne, BME001-05) in six replicates per condition. Organoids were cultured in Advanced DMEM/F-12 supplemented with 1x GlutaMAX, 10 mM HEPES, 1x Penicillin–Streptomycin, 1x B-27 supplement, 1.25 mM N-acetyl-L-cysteine (Sigma, A9165), 5 ng/mL EGF, 100 ng/mL Noggin, 500 ng/mL R-spondin (Thermo Fisher, 120-38), 500 nM A83-01 (Tocris Bioscience 2939), 10 ng/mL FGF-10 (Thermo Fisher, 100-26), 5 ng/mL FGF-2 (Thermo Fisher, 100-18B), 10 mM Nicotinamide (Sigma, N0636), 37.5 ng/mL Heregulin β-1 (Thermo Fisher, 100-03), 10 μM Y-27632 (Abmole Bioscience, M1817). Cells were treated with HHT (25, 100 nM) and Trametinib (1, 10 nM) either alone or in combination starting 24 h after plating. Organoids were imaged (Fig.S10B) on Day 1 and at treatment endpoint (Day 5 for PDX-267, and Day 7 for PDX-327 and PDX-365) followed by viability quantification (Fig.7B).

### Method details

#### Viability

Viability was assessed using CellTiter-Glo 3D (Promega, G9681) according to the manufacturer’s instructions. Luminescence quantification was performed using the Cytation 5 Multi-Mode Reader (Agilent) or the SpectraMax MiniMax 300 Cytometer (Molecular Devices).

#### Brightfield imaging

Murine organoids were imaged at 2.5x magnification using the Cytation 5 Multi-Mode Reader. Objects were masked using the cyto3 model of Cellpose(79), counted, and areas were quantified in ImageJ. Objects with an area below 50 μm^2^ or touching the well edges were excluded from quantification. Organoid phenotypes were quantified per brightfield image (triplicate per genetic/condition), with all objects ≥50 μm^2^ categorised as cystic or hyperplastic. The cystic cutoff threshold varied based on lumen thickness and organoid circularity. PDX-O Imaging was performed on the Opera Phenix Plus (Revvity). In each well, image stacks of 26.5 µm between planes over 1.4 mm were acquired using a 5x air objective across a 3x3 field of view. Objects were masked and area quantified using the Harmony software (Revvity).

#### Western blot

After 5 days of culture, Matrigel was removed with TrypLE (Thermo Fisher, 12605010) and organoids were lysed in NP-40 buffer (Thermo Fisher, J60766.AP) supplemented with protease (Sigma, 04693132001) and phosphatase inhibitor cocktails (Thermo Fisher, 78420). Proteins were separated by SDS–PAGE followed by western blotting and chemiluminescent detection using anti-mouse and anti-rabbit HRP conjugated secondary antibodies (Cell Signalling Technology, 7074 and 7076) (Fig.S1C).

#### Bulk RNA sequencing

RNA from organoids and minced tumours was extracted from TRIzol (Thermo Fisher, 15596026) lysates. Briefly, 1 volume of chloroform was added per 5 volumes of TRIzol. Samples were mixed and incubated 3 min, after which the upper phase was transferred to a PureLink RNA mini column (Thermo Fisher, 12183018A). RNA was extracted following the manufacturer’s instructions. Samples were treated with PureLink DNase (Thermo Fisher, 12185010) and eluted in ultrapure water. Library preparation and sequencing were performed by Azenta Life Sciences (Germany). Sequencing depth was set to 20 million paired-end reads per sample.

#### Reverse transcription and quantitative PCR

Reverse-transcribed cDNA was generated using RevertAid First Strand cDNA Synthesis Kit (Thermo Fisher, K1622) following the manufacturer’s instructions and amplified using gene-specific primers (Key Resources Table). Expression levels were normalised to TBP.

#### Organoid tissue microarray (TMA)

Murine organoids were fixed *in situ* using 4% paraformaldehyde (PFA) at 4°C overnight. Following removal of Matrigel through PBS washes, samples were resuspended in Histogel (Thermo Fisher, 12006679) and transferred to Tissue-Tek cryomolds (10 × 10 × 5 mm). Once solidified, samples were placed in cassettes and fixed overnight in 10% formalin at 4°C. Following paraffin embedding, TMA cores were extracted in triplicate at the Francis Crick Histology Core Facility using the TMA Master II (3DHISTECH).

#### H&E staining

Paraffin embedding and sectioning were performed by the Histopathology Core Facility at the Institute of Cancer Research.

#### Organoid TMA

Sections (2–3 μm) were dewaxed and hydrated, incubated for 7 min in Modified Harris haematoxylin solution (TCS Biosciences, HS351-1), differentiated in 1% acid alcohol and blued in Scott’s Tap Water Substitute (Leica, 3802900). Sections were then incubated in 1% Eosin Y (TCS Biosciences, 5091), washed in tap water, dehydrated, cleared in xylene (Fisher Scientific, 10385910), and mounted in DPX mountant (Sigma, 06522). Imaging was performed using the VS200 Research Slide Scanner (Olympus).

#### Tumour samples

Tumours were fixed in PFA 4% overnight and paraffin embedded. Sections (5 μm) were dewaxed and dehydrated through two changes of xylene and graded alcohols, rinsed in water, and nuclei stained with Harris Hematoxylin Nuclear Stains (Leica Biosystems, 3801560). Slides were differentiated in 1% acid alcohol, blued in running tap water, counterstained with 1% eosin Y, dehydrated through graded alcohols, cleared in xylene, and coverslipped. Imaging was performed using the NanoZoomer-XR C12000 (Hamamatsu).

#### Immunohistochemistry (IHC)

IHC staining was performed by the Histopathology Core Facility at the Institute of Cancer Research. Briefly, following tissue rehydration, antigen retrieval was performed using the Autostainer Link 48 (Agilent) at 97°C for 20 min. Endogenous peroxidases were blocked for 10 min with Dako REAL peroxidase-blocking solution (Agilent, S202386-2). Primary antibodies were incubated for 1 h at room temperature, secondary antibodies were incubated for 20–30 min at room temperature. Anti-rabbit ImmPRESS polymer-HRP (Vector Laboratories MP-7401) was used for detection. Images were captured using the NanoZoomer-XR C12000 (Hamamatsu).

#### Cyclic immunofluorescence (CyCIF)

CyCIF staining was adapted from published protocols(80,81). Briefly, paraffin sections were rehydrated, and antigen retrieval was performed using 1X IHC Antigen Retrieval Solution - High pH (Thermo Fisher, 00-4956-58) at 95°C for 20 min. Endogenous peroxidases were blocked with hydrogen peroxide 4.5% in PBS supplemented with 20 mM NaOH for 45 min, followed by standard blocking with sheep serum (Abcam, ab7489) supplemented with Bovine Serum Albumin (BSA: Sigma, A7906). All primary antibodies were incubated at 1:100 for 1 h at room temperature, secondary antibodies were incubated at 1:400 for 20 min at room temperature (see Key Resource Table). Following image capture, fluorophores were bleached in PBS containing 4.5% hydrogen peroxide and 20 mM NaOH under bright light for 45 min. Images were captured using the LSM980 Airyscan 2 (Zeiss) for organoids and the Axio Scan Z1 (Zeiss) for tumour sections.

#### Organoid CyTOF sample preparation

The protocol for CyTOF sample preparation was adapted from Sufi *et al*.(82). Briefly, organoids were labelled with 25 μM ¹²⁷IdU (Standard BioTools, 201127) in culture for 30 min. Protease (Sigma, P8340) and phosphatase inhibitors cocktails (Sigma, 4906845001) were added during the last 5 min of incubation. Organoids were then fixed with 4% PFA for 1 h at 37°C, washed twice with PBS, and incubated with 250 nM ¹⁹⁴Cisplatin (Standard BioTools, 201194) in PBS for 10 min at room temperature on a rocker. Cells were washed twice with PBS then TOBis barcoding reagents (126-plex) were added to each well and incubated overnight at 4°C. Excess barcodes were quenched the next day with 2 mM L-Glutathione reduced (Sigma, G6529) in PBS for 10 min at room temperature on a rocker then washed twice with PBS. Samples were pooled and dissociated using a gentleMACS Octo Dissociator with heaters (Miltenyi Biotec 130-096-427; Loop: 1,500 rpm fwd spin for 2s, 1,500 rpm rev spin for 2s, 50 rpm fwd spin for 3 min at 37°C for 1 h) in a solution composed of DPBS with calcium and magnesium (Thermo Fisher, 11570456) supplemented with 2 mM MgCl₂, 2.5 mg/mL Dispase II (Thermo Fisher, 17105041), 0.2 mg/mL Collagenase IV (Thermo Fisher, 17104019), and 0.2 mg/mL DNase I (Sigma, DN25). Cells were washed with PBS and incubated for 15 min at 37°C in 0.2 mg/mL DNase I in DPBS with calcium and magnesium supplemented with 2 mM MgCl₂, followed by a wash in PBS containing 2 mM EDTA. Cells were filtered (40 μm), counted then stained with a panel of metal-conjugated extracellular antibodies (Table S5) in Cell Staining Buffer (CSB: Standard BioTools, 201068) supplemented with TruStain FcX (BioLegend, 101320) at room temperature for 30 min on a rocker.

Cells were then washed in CSB and fixed with 4% PFA for 20 min at room temperature, then permeabilised with FoxP3 Fix/Perm Buffer (Fisher Scientific, 11500597) for 30 min at 4°C, followed by 50% methanol for 10 min on ice. After washing twice with CSB cells were stained for intracellular and phospho-proteins in CSB supplemented with TruStain FcX (BioLegend, 101320) (Table S5) for 30 min at 4°C on a rocker. Cells were washed in CSB then fixed in 1.6% formaldehyde for 10 min. Cells were stored in Fix/Perm buffer (Standard BioTools, 201067) containing 1 μM ¹⁹¹/¹⁹³Ir DNA intercalator (Standard BioTools, 201192A) at 4°C overnight before aquisition.

#### Immune CyTOF sample preparation

Collected tumours were minced and dissociated using the Tumour Dissociation Kit (Miltenyi Biotec, 130-096-730) according to the manufacturer’s protocol. Cells were washed with PBS, then incubated in ACK Lysis Buffer (Thermo Fisher, A1049201) on ice for 1 min to remove red blood cells then neutralised with excess PBS. Cells were filtered and incubated with 1 μM ¹⁹⁶/¹⁹⁸Cisplatin (Standard BioTools 201196/8) in PBS for 10 min, washed twice, then incubated for 10 min at room temperature in CSB supplemented with TruStain FcX. Cells were then stained with metal-conjugated extracellular antibodies in CSB (Table S5) for 30 min at room temperature on a rocker. Cells were then washed with CSB and barcoded using the Cell-ID 20-Plex Pd Barcoding Kit (Standard BioTools 201060) following the manufacturer’s instructions, washed twice and pooled. Samples were fixed in 1.6% formaldehyde for 10 min at room temperature, then stored in Fix/Perm buffer containing 1 μM ¹⁹¹/¹⁹³Ir DNA intercalator at 4°C overnight. The next day, samples were frozen and stored at -80°C until acquisition.

#### CyTOF acquisition

Samples were washed in CSB supplemented with 2 mM EDTA and further washed with CAS solution (Standard BioTools, 201237). Cells were spiked with EQ Four Element Calibration Beads (Standard BioTools, 201078), diluted 1:10 using CAS solution with 2 mM EDTA, and acquired using a Helios Mass Cytometer (Standard BioTools).

#### Preclinical trial

Dissociated PtPM organoids were injected as described above. Tumours were monitored for 25 days, at which time all mice were chemically castrated via subcutaneous administration of Degarelix acetate (0.315 mg in 100 μL water; Cayman Chemical, 24069), repeated every 2 weeks. Mice were randomised to control or treatment arms, and drug administration began 48 h after castration. All compounds were administered once daily for 5 days, followed by 2 days off over a 25 - day time course. Trametinib and Afatinib were dosed PO at 1mg/kg and 10mg/kg, respectively. Both compounds were administered in a vehicle of PBS, 0.5 (w/v) % Methylcellulose (Sigma, M6385), and 0.4 (v/v) % Tween-80 (Sigma, P1754). HHT was dosed 1.0mg/kg IP in PBS. Control mice were dosed in accordance with the combination therapy. Weights were monitored 5 days a week, and mice were euthanised when tumour volume approached 1.5 cm³.

#### Oncoprint alteration frequency analysis

Genomic alteration data were downloaded from cBioPortal (MSK/DFCI 2018, MSK 2022, MSK 2020, SU2C 2015, SU2C 2019, see Key Resources Table). Gene-level aberrations in TP53, ERG, PTEN, MYC, APC and RB1 were annotated as deep deletions, amplifications, structural variants, or putative driver/passenger mutations, with copy-number changes taking precedence over SVs and mutations. Samples were grouped by study, stratified by metastatic status, and ordered hierarchically by alteration type (deep deletion > amplification > SV > driver > passenger > none). Oncoprints were generated using Python packages *pandas (*https://github.com/pandas-dev/pandas) and *matplotlib* (https://github.com/matplotlib/matplotlib) with distinct colour codes for each alteration and study. Aberration frequencies were calculated for all samples, by study, and stratified by metastatic status. Co-occurrence and mutual exclusivity between gene pairs were tested by Fisher’s exact test.

### Quantification and statistical analysis

#### Mass cytometry data processing

Raw CyTOF data were normalised and exported as standard FCS files. Multiplexed TOB *is* and Cell-ID labelled experiments were de-barcoded to separate experimental conditions (https://github.com/zunderlab/single-cell-debarcoder). Gating with Gaussian parameters and Ir intensity removed debris and doublets, followed by cisplatin for live-dead separation. Processing adapted from Qin *et al*.(63).

#### Mass cytometry analysis

Earth Mover’s Distance (EMD) scores were calculated using the CyGNAL 0.2.1 pipeline(82). Heatmaps were generated from mean EMD values using the *ggplot2*(83) and *tidyverse*(84) R packages. PHATE embedding was computed from EMD scores using the *PHATE* package for R(85) (*k*NN = 5, T = 5).

#### RNA sequencing

##### Preprocessing

Raw sequencing reads were processed with the nf-core RNA-seq pipeline(86) using Nextflow. Reads were aligned to the GRCm38 (mm10) reference genome and Ensembl v95 gene annotation using the “star_salmon” aligner with default options. Compiled raw counts were filtered (genes were kept if at least 10% of samples had at least 1 transcript per million), and *ComBat-seq*(87) R package was run with Type (i.e. Organoid/Allograft) as the ‘batch’. Rounded counts were fed into *DESeq2* (v1.48.2)(88) and variance-stabilising transformed (VST) counts were used for downstream analysis.

##### Differential expression

Differential expression analyses were completed with *DESeq2* and GSEA with *clusterProfiler*(89) R packages. The pathway databases explored were GO Biological Processes (90) and MSigDB Hallmark(91). Volcano plots were made using the *EnhancedVolcano* R package (https://github.com/kevinblighe/EnhancedVolcano). The R package *GSVA* (v2.2.0)(92) was used to get single sample scores of MSigDB Hallmark pathways. Heatmaps were made with the *pheatmap* R package (https://github.com/raivokolde/pheatmap).

##### Signatures

Publicly available datasets were downloaded (details in Key Resources Table and Table S3), converted to a common set of gene IDs using *biomaRt*(93) (v.2.64.0) and transformed into log2 (TPM+1). For organoid analysis, human Ensembl IDs were used (DepMap cell line data downloaded from https://depmap.org/portal/ already in correct units), while mouse Ensembl IDs were used for the allograft analysis. Non-normalised ssGSEA (via the *GSVA* R package) was run on a compiled list of signatures. Gene lists available in Table S2.

##### Transcriptional alignment and HuMo analysis

For TCGA data(94), gene counts were downloaded and compiled from the GDC web portal (https://portal.gdc.cancer.gov/), and *DESeq2* and VST were run. For SU2C(95), raw gene counts were estimated from available fragments per kilobase of transcript per million mapped reads (FPKM), assuming a total library size of ∼50 million reads and using gene lengths from the Gencode V26 GTF. These were then processed and normalised using *DESeq2* and VST. The resulting normalised data were scaled by row and column separately for ProMPt, TCGA, and SU2C cohorts. These cohorts were then combined, using *biomaRt* to convert mouse to human Ensembl gene IDs. The HuMo analysis method was then carried out as in Müller *et al.*(18) using TCGA & SU2C as the human sample to map the mouse data onto.

Lastly, Han *et al*.(11) and Tang *et al*.(9) signature scores were determined using ssGSEA (from the *GSVA* R package) on the scaled expression data of the HuMo analysis. A permutation analysis was then performed to determine if any signatures were significant on a sample-by-sample basis. For Han, if only one signature was significant, the sample was assigned to that subtype, but if none or multiple signatures were significant, the sample was assigned to ‘Mixed’. For Tang, the sample was assigned to the subtype with the smallest p-value regardless of significance (first alphabetically chosen in the rare occurrence of multiple p-values at the minimum).

##### Gene signature correlation analysis

Transcriptomic data from two mCRPC cohorts (SU2C–PCF(95) and RMH(96)) were downloaded and reanalysed. RNAseq reads were aligned to the human reference genome (GRCh37/hg19) using *TopHat2*(97) (v.2.0.7). Gene expression was quantified as FPKM using *Cufflinks*(98) (v.2.2.1). Hallmark and Reactome signatures were obtained from MSigDB (v7.1). Pathway score was computed with GSVA, and associations between signatures were analysed using the two-sided Spearman’s rank correlation test.

##### Statistical analysis

All statistical details, including the statistical tests used, the number of samples, and output, can be found in Figures, Figure legends, or Results. Data are presented as mean ± SD or ± SEM with individual biological replicates shown. For *in vivo* experiments, each subcutaneous tumour was considered an independent sample.

Analyses were performed using GraphPad Prism or in RStudio. The R package *SQUEAK* (https://github.com/seedgeorge/SQUEAK) was used to achieve mixed-effect regression modelling and visualisation.

## KEY RESOURCES TABLE

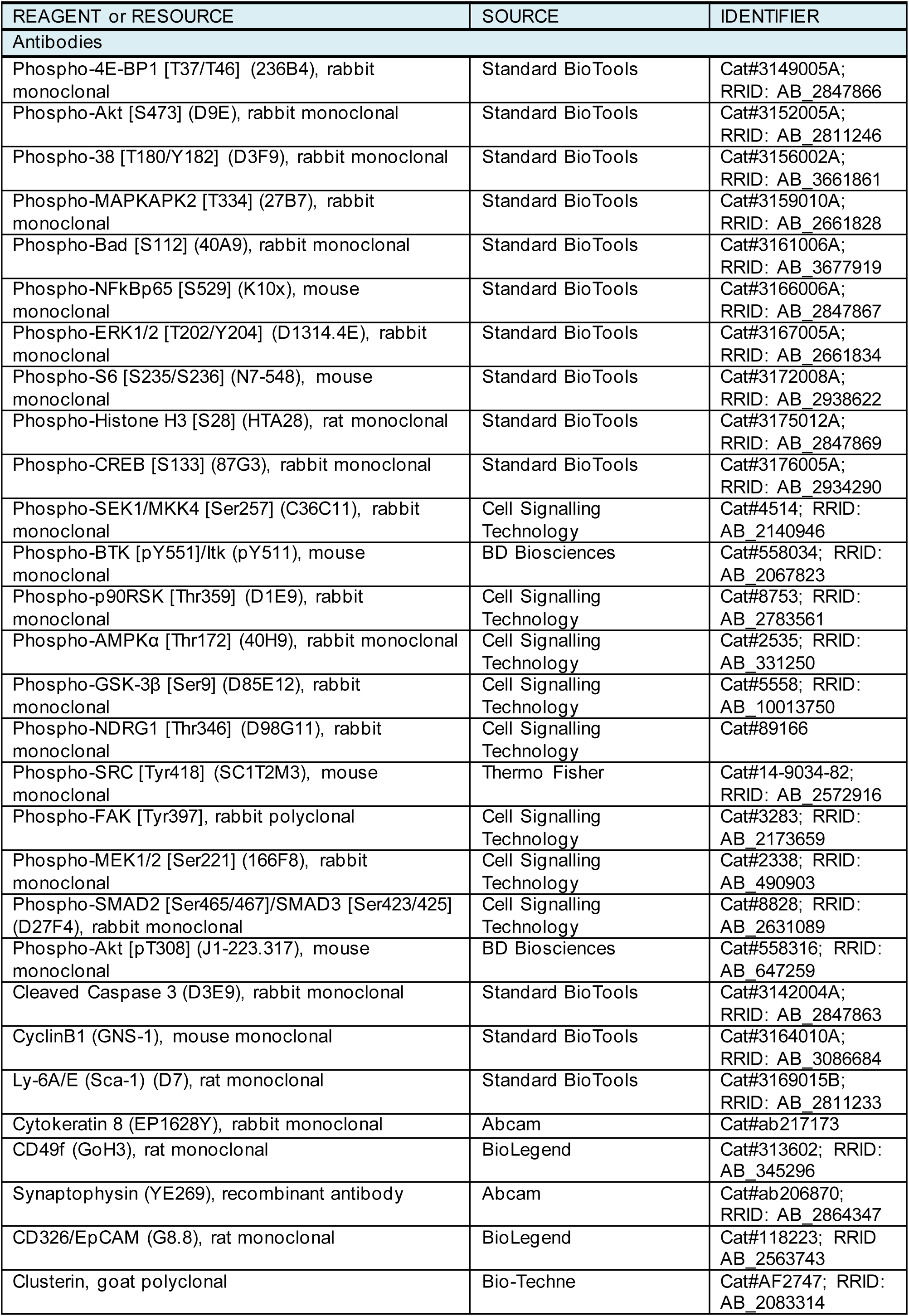
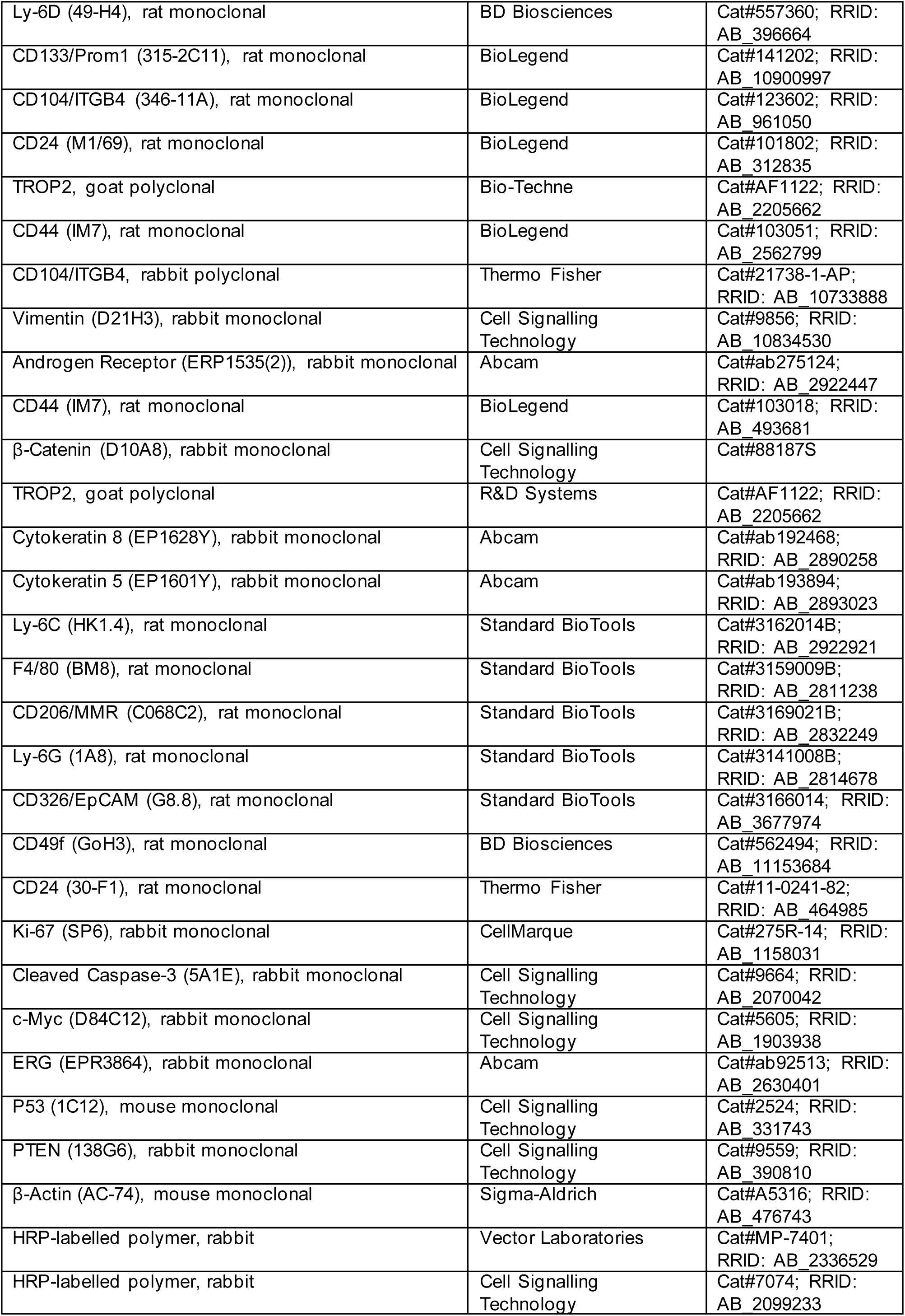
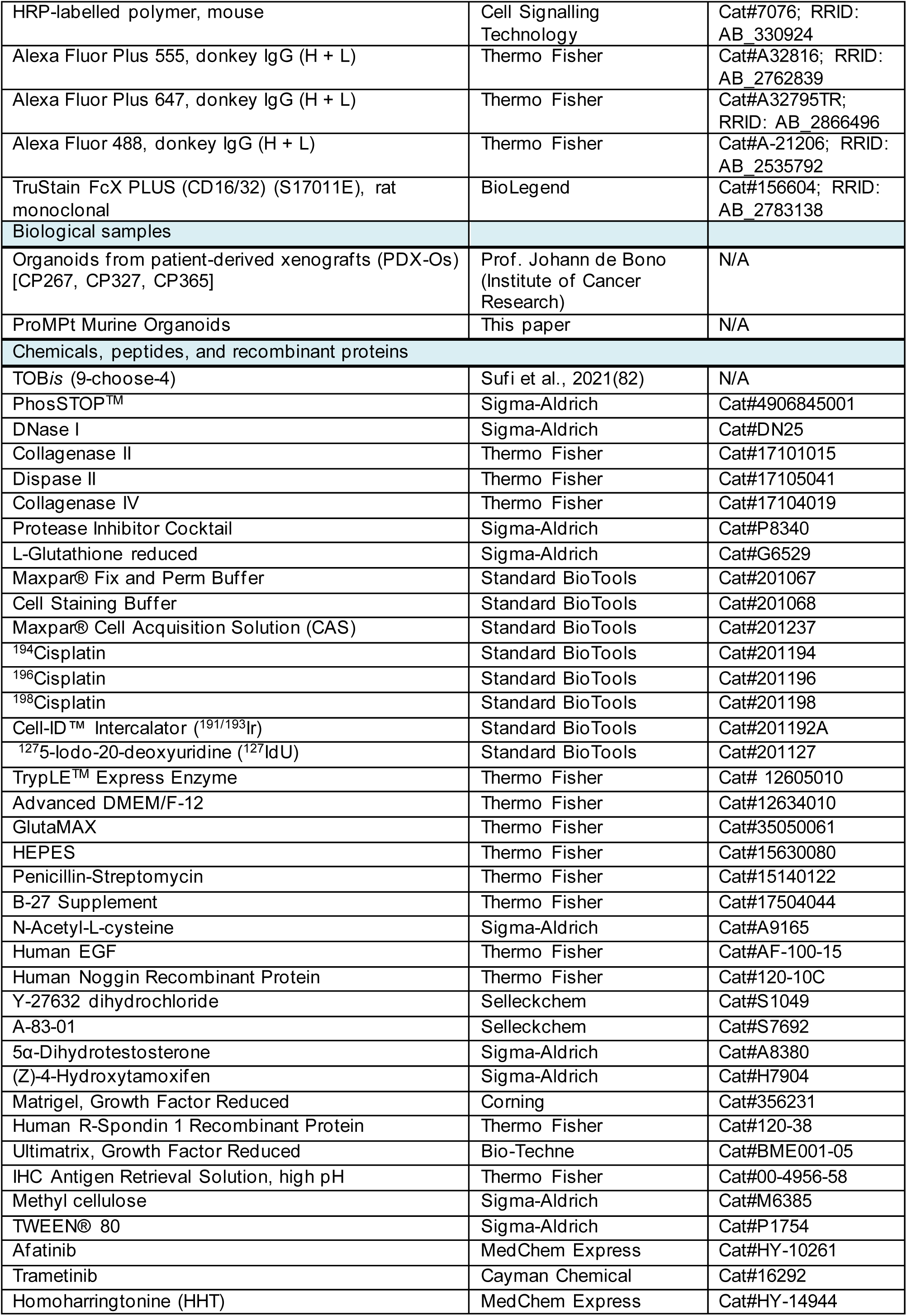
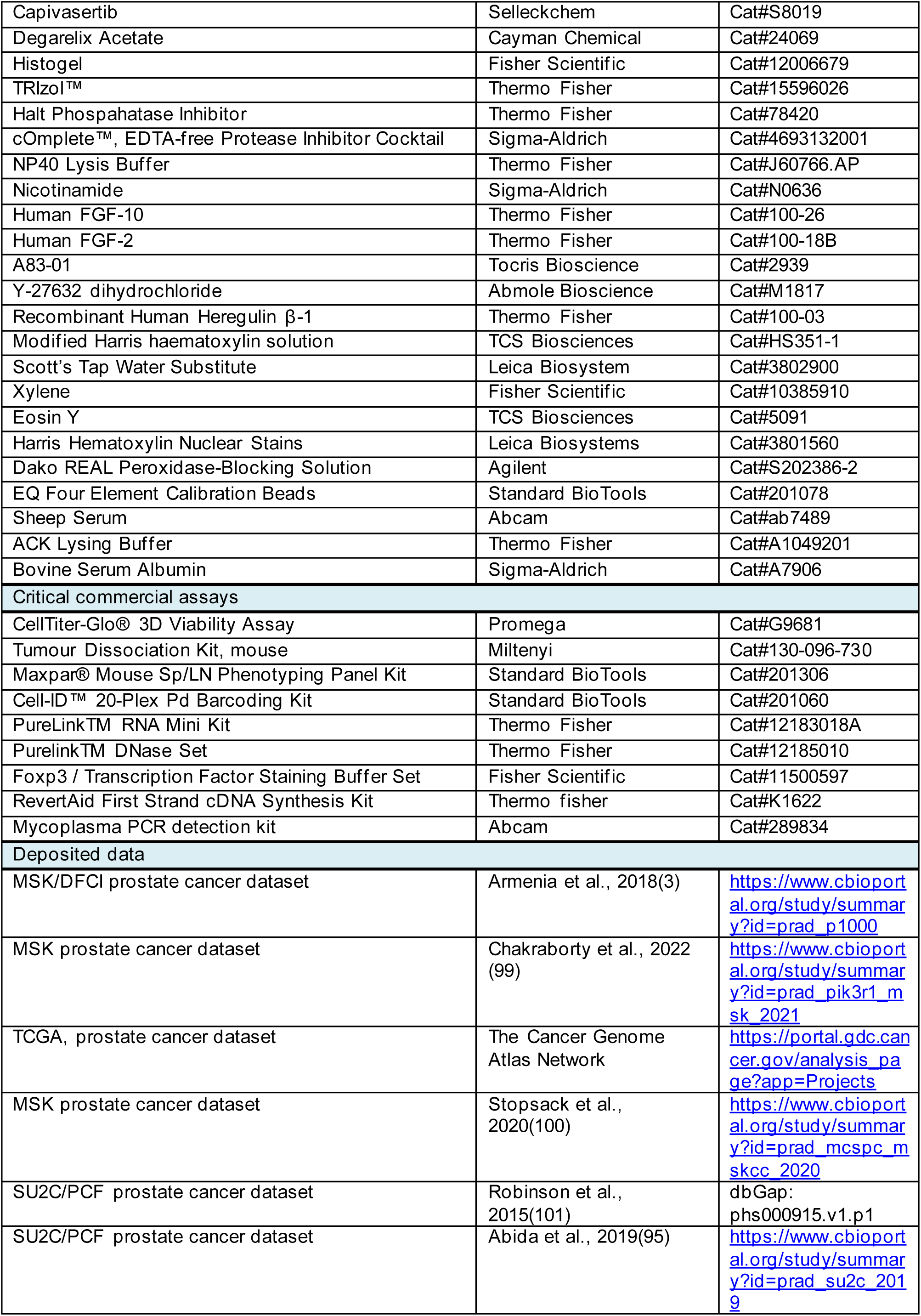
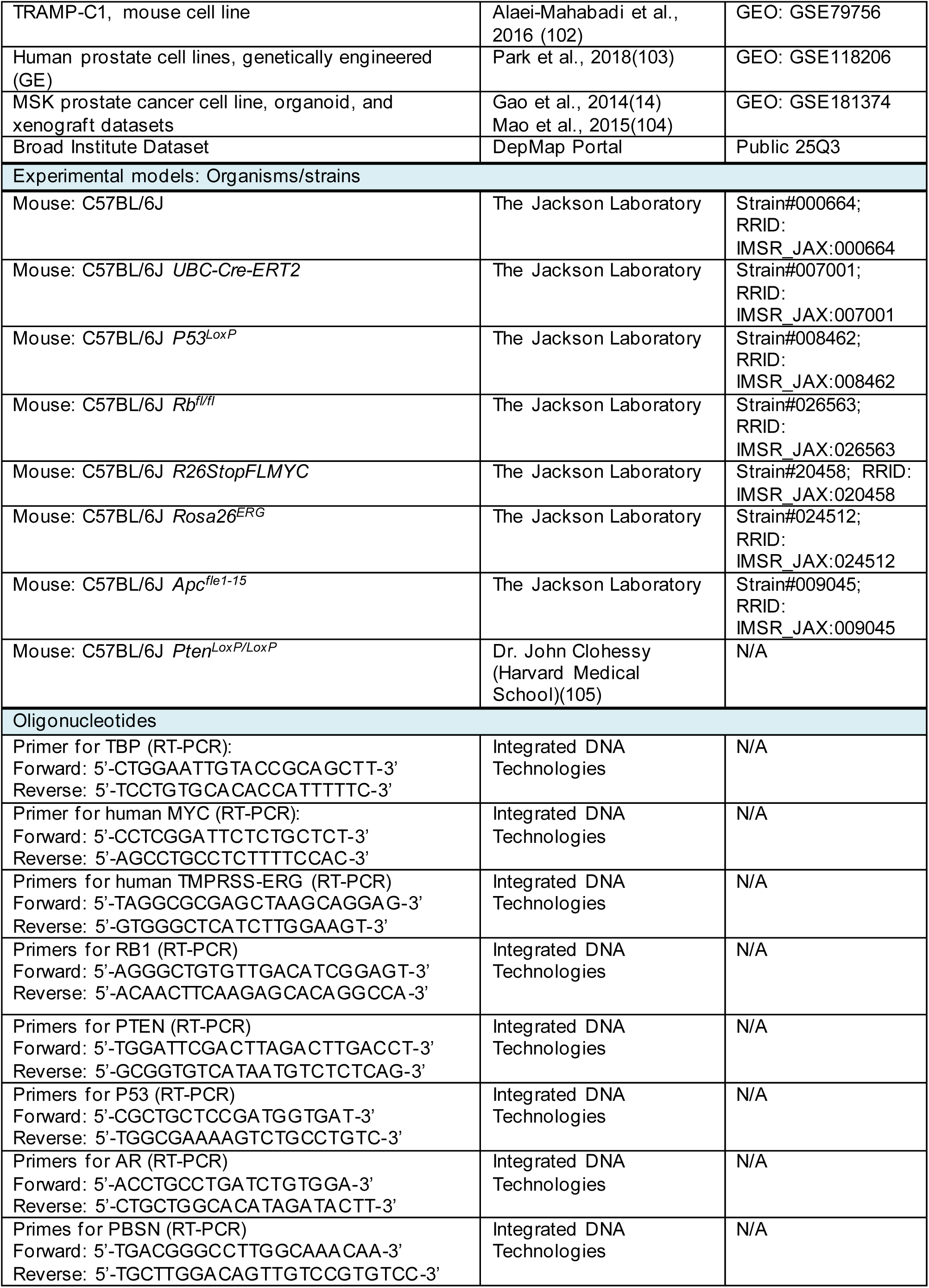
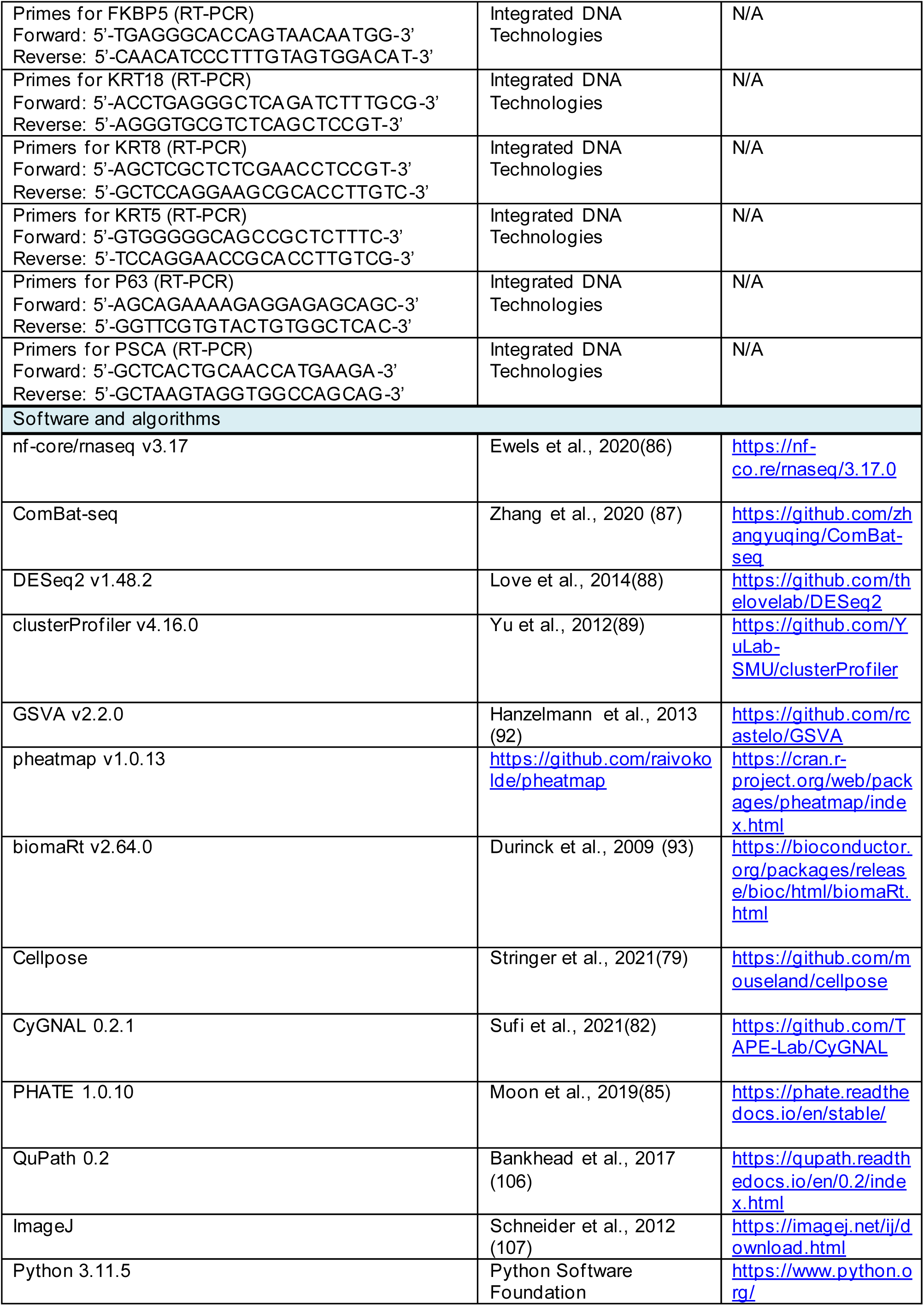
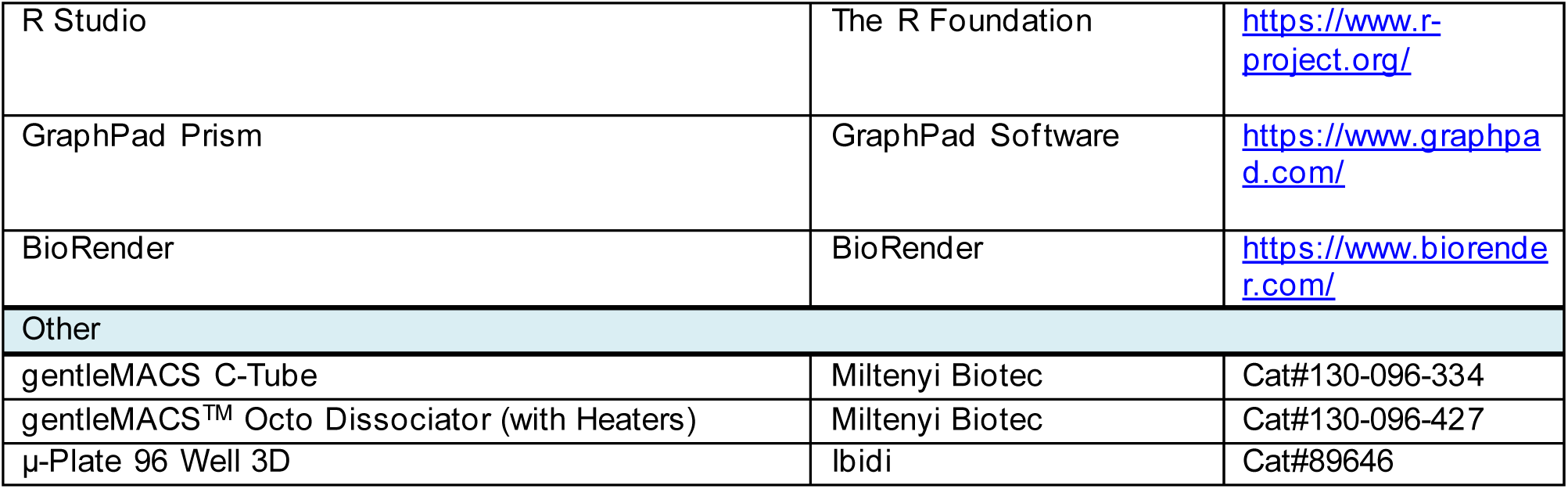

## Data availability statement

The sequence data and the mitochondrial and chloroplast sequences generated or analyzed in this study are available in GenBank (PX438801, PV798382, and PV826156), the Sequence Read Archive (SRA: SRR31014706, SRR31014705).

## Supplemental Information

Document S1: Supplementary Figures 1-11

Supplementary table 1: Histopathological assessment of ProMPt tumours

Supplementary table 2: Gene signatures used in data analysis. Related to Figure 2, S2, 5, S8.

Supplementary table 3: List of publicly available models used in signature assessment. Related to Figure 2, and S2

Document S2: Supplementary Table 4, Genomic details of PDX-O samples. Related to Figure 7 and S10; Supplementary Table 5, CyTOF Antibody Panels. Related Figures 4, 6, 7, and Figure S6, S7, S9, and S11.

