## Supplementary material for "ProMPt: A modular preclinical platform for functional modelling of prostate cancer heterogeneity and therapeutic vulnerabilities": Document S1 - Supp Figures 1-11

### Supp Fig. 1

## A

| Aberrant per Study | % aberrant samples (all 6 genes) | TP53 | ERG | PTEN | MYC | APC | RB1 |
|---|---|---|---|---|---|---|---|
| Prostate Adenocarcinoma (MSK/DFCI, Nature Genetics 2018) | 60.320 | 24.82 | 32.30 | 20.02 | 7.83 | 5.52 | 6.14 |
| Prostate Adenocarcinoma (MSK, Clin Cancer Res. 2022) | 66.289 | 31.38 | 25.95 | 24.52 | 8.71 | 11.22 | 6.26 |
| Metastatic castration-sensitive prostate cancer (MSK, Clin Cancer Res 2020) | 62.424 | 35.76 | 25.45 | 18.18 | 6.06 | 9.70 | 6.06 |
| Metastatic Prostate Cancer (SU2C/PCF Dream Team, Cell 2015) | 85.938 | 42.19 | 48.44 | 39.06 | 23.44 | 14.06 | 9.38 |
| Metastatic Prostate Adenocarcinoma (SU2C/PCF Dream Team, PNAS 2019) | 77.723 | 41.09 | 30.94 | 32.67 | 24.75 | 7.67 | 13.61 |
| All Samples | 2261 / 3433 (65.9%) | 30.79 | 29.01 | 23.97 | 10.46 | 8.91 | 7.14 |
| Prostate only | 1240 / 2070 (59.9%) | 26.18 | 30.29 | 18.55 | 6.47 | 6.28 | 4.25 |
| Other Tissues only | 1021 / 1363 (74.9%) | 37.78 | 27.07 | 32.21 | 16.51 | 12.91 | 11.52 |

| Samples | Gene1 | Gene2 | Both | G1_only | G2_only | Neither | N | Cooccur_% | OddsRatio | Fisher_p(two-sided) | Direction |
|---|---|---|---|---|---|---|---|---|---|---|---|
| All | TP53 | ERG | 406 | 651 | 590 | 1786 | 3433 | 11.83 | 1.89 | 2.80E-12 | co-occur |
| All | TP53 | PTEN | 367 | 690 | 456 | 1920 | 3433 | 10.69 | 2.24 | 2.57E-12 | co-occur |
| All | ERG | PTEN | 347 | 649 | 476 | 1961 | 3433 | 10.11 | 2.20 | 2.78E-12 | co-occur |
| All | TP53 | MYC | 163 | 894 | 196 | 2180 | 3433 | 4.75 | 2.03 | 7.81E-10 | co-occur |
| All | TP53 | RB1 | 130 | 927 | 115 | 2261 | 3433 | 3.79 | 2.76 | 1.34E-12 | co-occur |
| All | PTEN | RB1 | 103 | 720 | 142 | 2468 | 3433 | 3.00 | 2.49 | 9.41E-11 | co-occur |
| Prostate | TP53 | ERG | 214 | 328 | 413 | 1115 | 2070 | 10.34 | 1.76 | 9.17E-08 | co-occur |
| Prostate | ERG | PTEN | 171 | 456 | 213 | 1230 | 2070 | 8.26 | 2.17 | 7.38E-11 | co-occur |
| Prostate | TP53 | PTEN | 165 | 377 | 219 | 1309 | 2070 | 7.97 | 2.62 | 1.36E-12 | co-occur |
| Prostate | TP53 | MYC | 56 | 486 | 78 | 1450 | 2070 | 2.71 | 2.14 | 6.12E-05 | co-occur |
| Prostate | TP53 | RB1 | 42 | 500 | 46 | 1482 | 2070 | 2.03 | 2.71 | 9.72E-06 | co-occur |
| Prostate | TP53 | APC | 39 | 503 | 91 | 1437 | 2070 | 1.88 | 1.22 | 0.304139845 | co-occur |
| Other Tissues | TP53 | PTEN | 202 | 313 | 237 | 611 | 1363 | 14.82 | 1.66 | 2.12E-05 | co-occur |
| Other Tissues | TP53 | ERG | 192 | 323 | 177 | 671 | 1363 | 14.09 | 2.25 | 7.36E-11 | co-occur |
| Other Tissues | ERG | PTEN | 176 | 193 | 263 | 731 | 1363 | 12.91 | 2.53 | 1.96E-12 | co-occur |
| Other Tissues | TP53 | MYC | 107 | 408 | 118 | 730 | 1363 | 7.85 | 1.62 | 0.001184676 | co-occur |
| Other Tissues | TP53 | RB1 | 88 | 427 | 69 | 779 | 1363 | 6.46 | 2.33 | 1.15E-06 | co-occur |
| Other Tissues | PTEN | RB1 | 70 | 369 | 87 | 837 | 1363 | 5.14 | 1.83 | 0.000547994 | co-occur |

## B

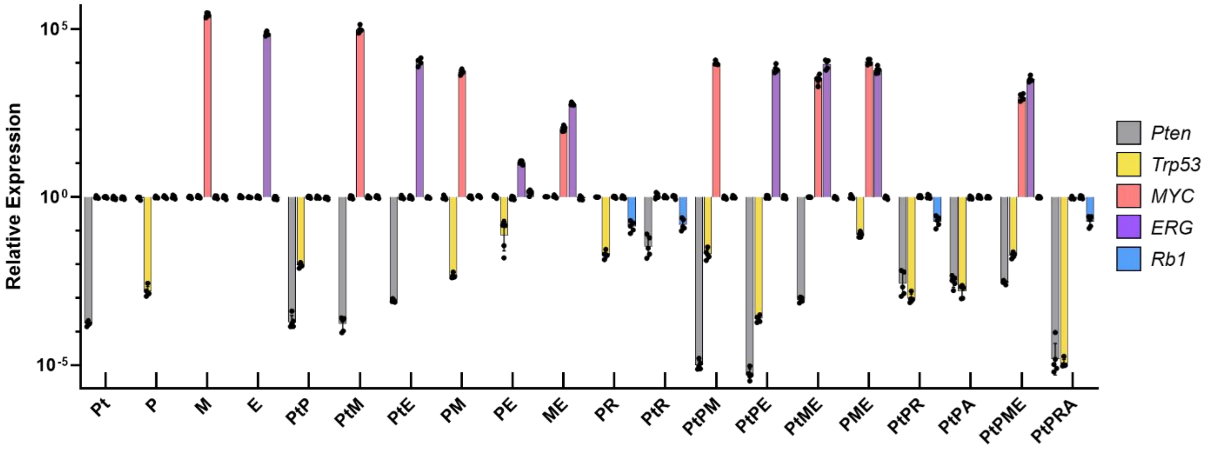


## C

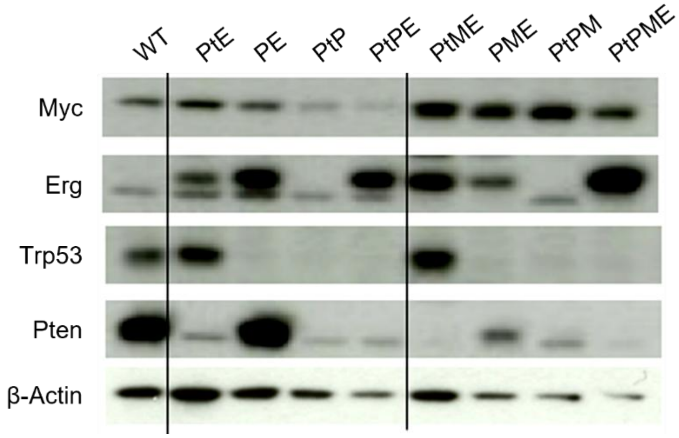


## D

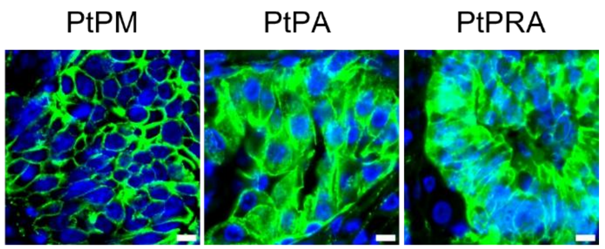

**Supplementary Figure 1. ProMPt: An organoid pipeline to model frequent prostate cancer drivers.**

(A) Top: frequency of individual genetic alterations segmented by study or disease site. Bottom: top five co-occurring gene pairs ranked by frequency across cohorts and disease site.

(B) RT–PCR analysis of *Pten*, *Trp53*, *MYC*, *Erg*, and *Rb1* transcript levels in organoids following Cre-mediated recombination. Error bars indicate mean ± SD of technical replicates (n = 5 per genotype).

(C) Western blot analysis of MYC, Erg, Trp53, and Pten protein levels in 4-OHT treated ProMPt organoids.

(D) Representative immunofluorescence (IF) staining of β-catenin (green) to assess Apc deletion. Scale bar, 50 μm.

Supp Fig. 2

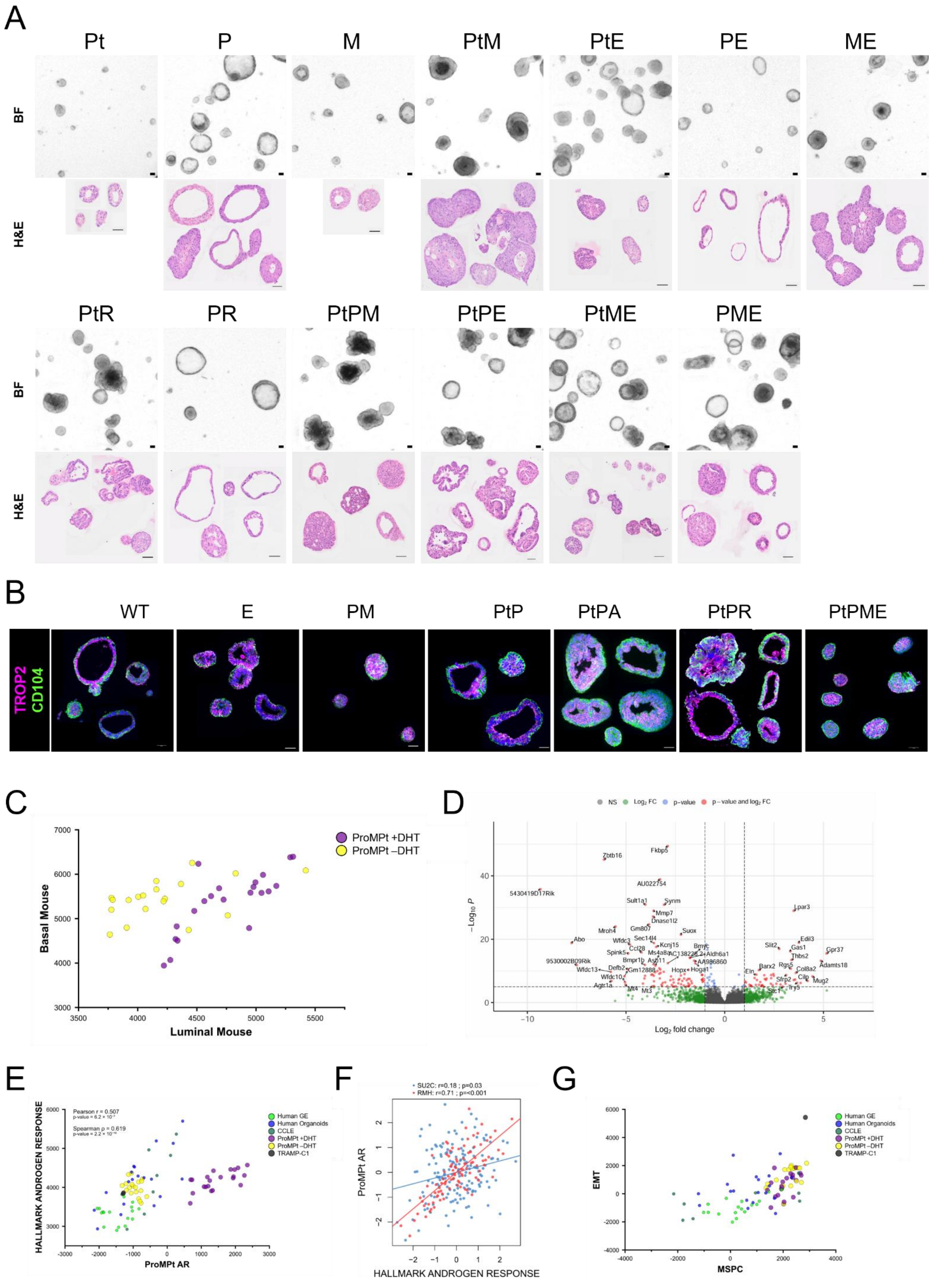
A
Pt
P
M
PtM
PtE
PE
ME
BF
H&E
PtR
PR
PtPM
PtPE
PtME
PME
B
WT
E
PM
PtP
PtPA
PtPR
PtPME
TROP2
CD104
C
ProMPt +DHT
ProMPt –DHT
Basal Mouse
Luminal Mouse
D
NS
p−value
p−value and log2 FC
Log2 fold change
−Log10 P
Fkbp5
Zbtb16
AU022754
5430419D17Rik
Sult1a1
Synm
Lpar3
Mmp7
Dnase1l2
Gm807
Suox
Mroh4
Abo
Wfdc3
Sec14l4
Kcnj15
Slit2
Edil3
Gas1
Gpr37
Thbs2
Adamts18
Rgs5
Col8a2
Barx2
Eln
Sfrp2
Cilp
Mug2
Tnfrsf1
Try5
Stc1
Bmyc
Aldh6a1
AA986860
Hopx
Hoga1
Ccl28
Spink5
Ms4a8a
Bmpr1b
Asb11
9530002B09Rik
Wfdc13
Defb2
Gm12888
Wfdc10
Agtr1a
Mt4
Mt3
E
HALLMARK ANDROGEN RESPONSE
ProMPt AR
Pearson r = 0.507
Spearman ρ = 0.619
Human GE
Human Organoids
CCLE
ProMPt +DHT
ProMPt −DHT
TRAMP-C1
F
SU2C: r=0.18 ; p=0.03
RMH: r=0.71 ; p=<0.001
ProMPt AR
HALLMARK ANDROGEN RESPONSE
G
EMT
MSPC

**Supplementary Figure 2. Characterization of ProMPt organoids following androgen withdrawal.**

(A) Representative brightfield (BF) and H&E images of ProMPt biobank organoids. Scale bars, 2 mm (BF) and 50 μm (H&E).

(B) Immunofluorescence (IF) staining of Trop2 and Cd104 in ProMPt organoid samples corresponding to those shown in Figure 2A. Scale bar, 50 μm.

(C) Scatterplot showing murine basal and luminal signature scores in ProMPt organoids under long-term DHT withdrawal compared with untreated conditions. Each dot represents one sample.

(D) Volcano plot showing $\log_2$ fold change in gene expression from bulk RNA-seq comparing untreated (n = 19) and long-term DHT-withdrawn (n = 17) ProMPt organoids. Genes meeting the indicated significance thresholds are colour coded according to the figure legend.

(E) Scatterplot showing expression of ProMPt-derived androgen receptor and HALLMARK_ANDROGEN_RESPONSE signatures across ProMPt organoids and established prostate cancer cell lines and organoids. Pearson's and Spearman's correlation coefficients are indicated. See also accompanying details in Supp. Table 3.

(F) Scatterplot showing the correlation between ProMPt Ar and HALLMARK_ANDROGEN_RESPONSE signatures in the SU2C (blue) and Royal Marsden Hospital (RMH; red) datasets. Pearson's correlation coefficients are indicated.

(E) Scatterplot showing expression of MSPC(10) and EMT signatures across ProMPt organoids and established prostate cancer cell lines and organoids. See also Supp. Table 3 for sample details and Supp. Table 2 for signature gene lists.

Supp Fig. 3

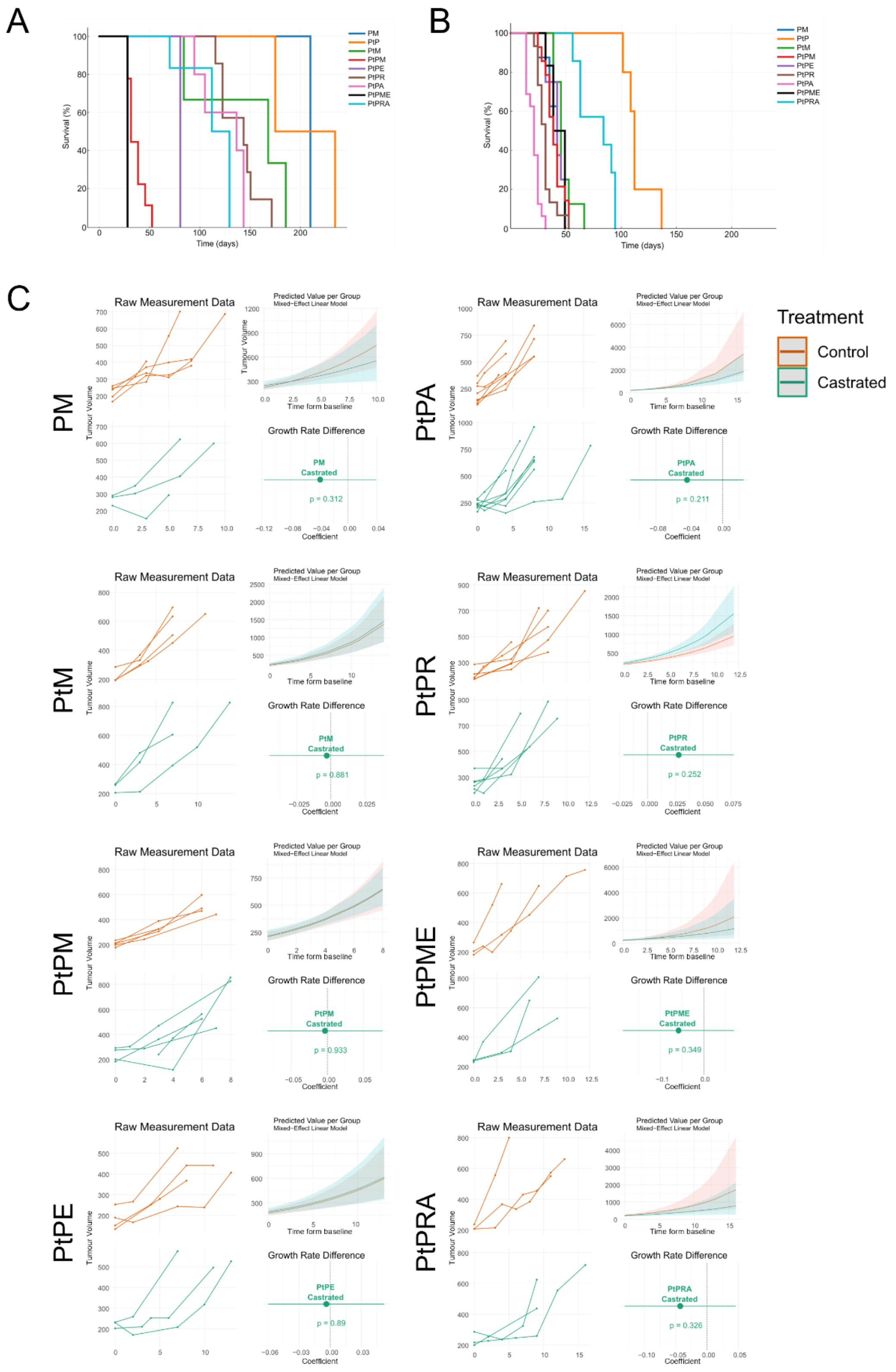


**Supplementary Figure 3. ProMPt tumours are castration resistant.**

(A) Kaplan–Meier survival curves for Gen. 1 ProMPt tumour models. Cohort sizes are indicated in Figure 3A.

(B) Kaplan–Meier survival curves for Gen. 2 ProMPt allograft models, including both intact and castrated cohorts, as castration did not significantly affect tumour growth or time to endpoint (see Figure S3C). Sample numbers are indicated in Figure 3A.

(C) Individual tumour growth curves of intact (orange) and castrated (green) mice across ProMPt allograft models. Each line represents one animal (minimum n = 3 per cohort). Predicted tumour growth was extrapolated from raw volume measurements using a linear mixed-effects model. Differences in growth rate between intact and castrated groups did not reach statistical significance across genotypes.

Supp Fig. 4

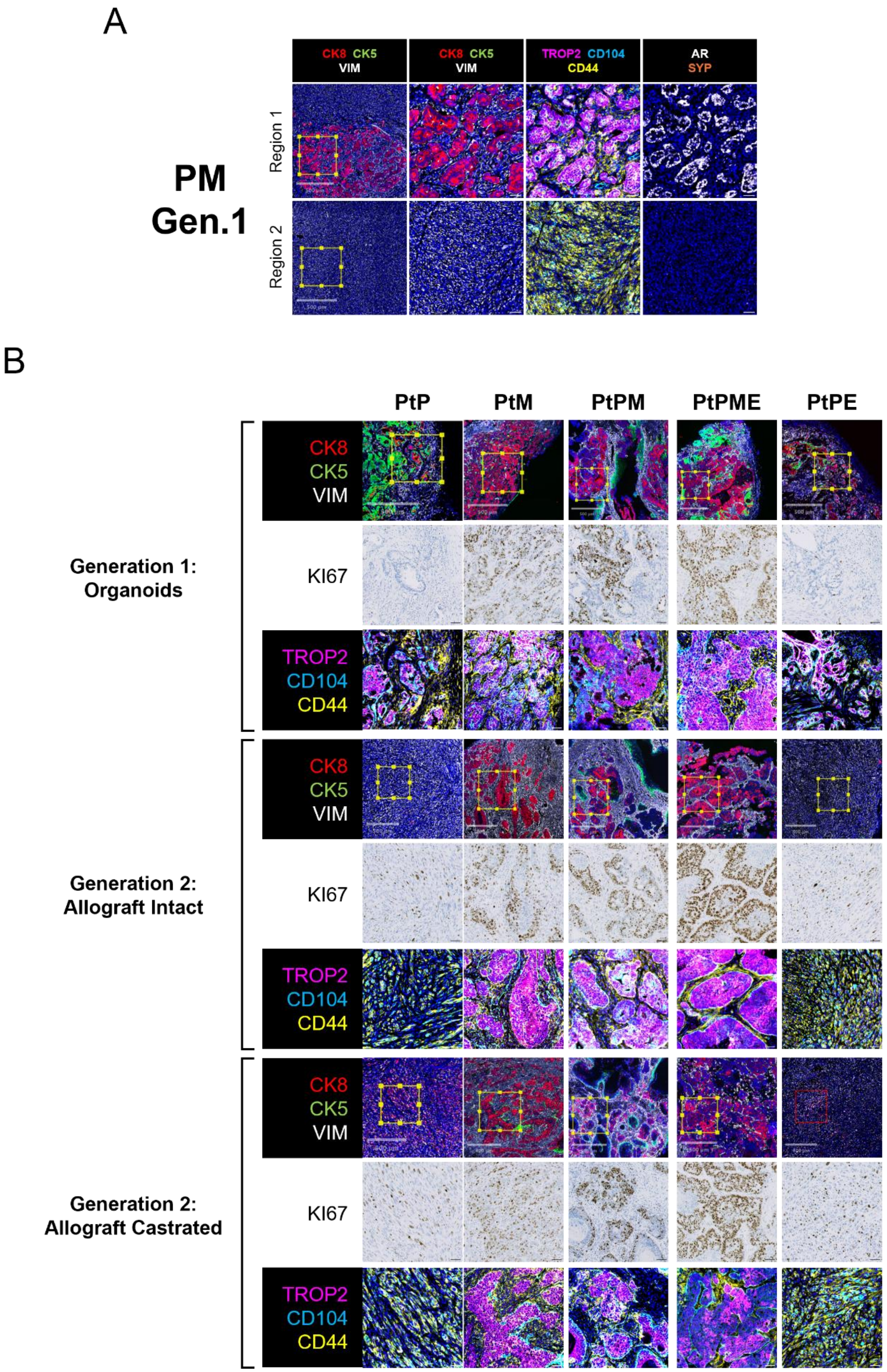


**Supplementary Figure 4. ProMPt tumours model frequent and rare CRPC subtypes.**

(A) Representative H&E and CyCIF images of Gen. 1 PM tumours stained for lineage and stem markers. Serial sections were taken between H&E and CyCIF. Scale bar, 50 µm.

(B) Regions of interest (ROIs) and additional immune staining for the indicated samples shown in Figure 3D. Scale bars, 500 µm (ROIs) and 50 µm (all staining).

Supp Fig. 5

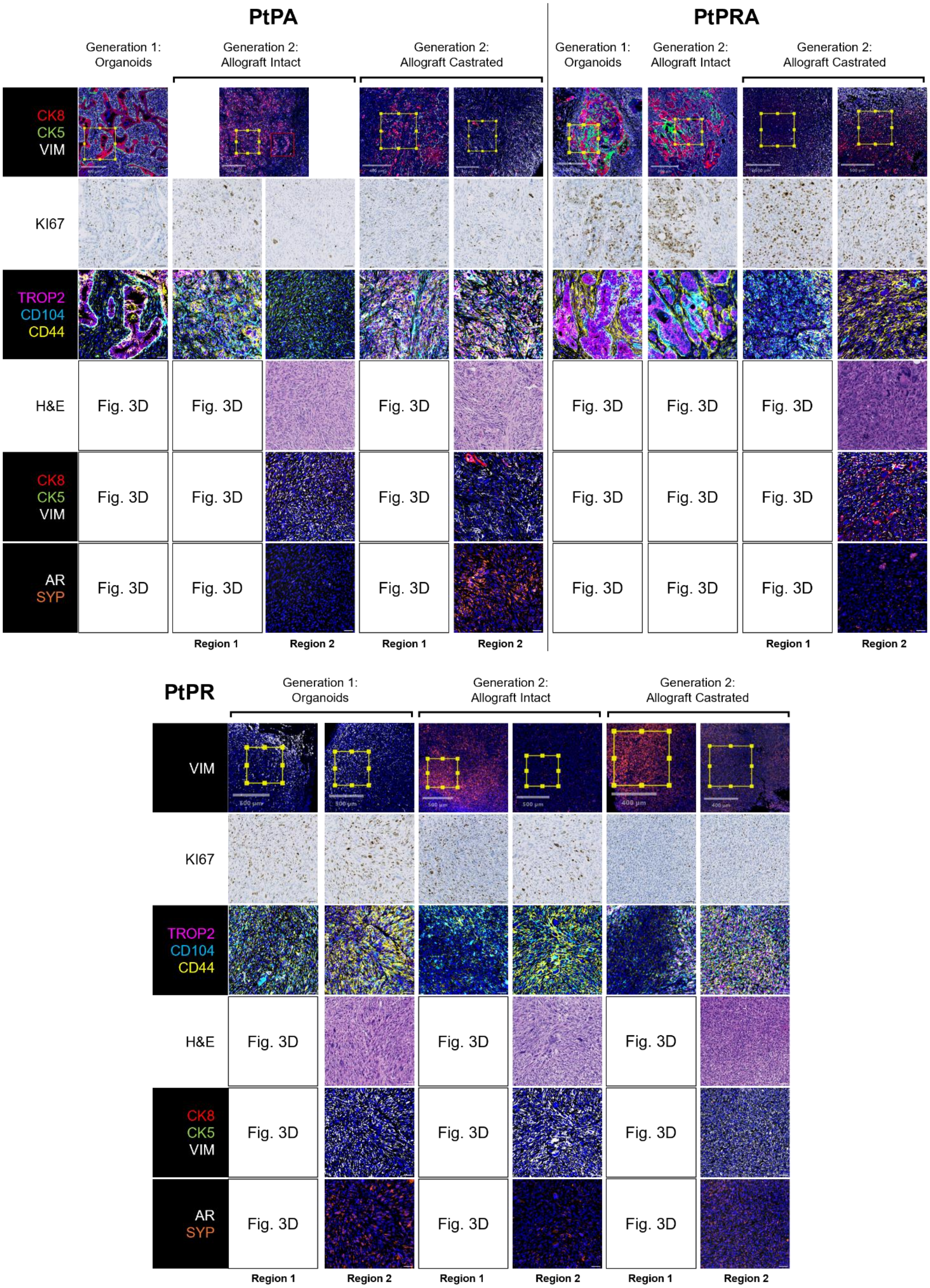


**Supplementary Figure 5. ProMPt tumours model frequent and rare CRPC subtypes.**

Regions of interest (ROIs) and additional immune staining for the indicated samples shown in Figure 3D, with full panels for alternative ROIs. Empty boxes denote images presented in the main-text Figure 3D. Scale bars, 500 µm (ROIs) and 50 µm (all staining).

### Supp Fig. 6

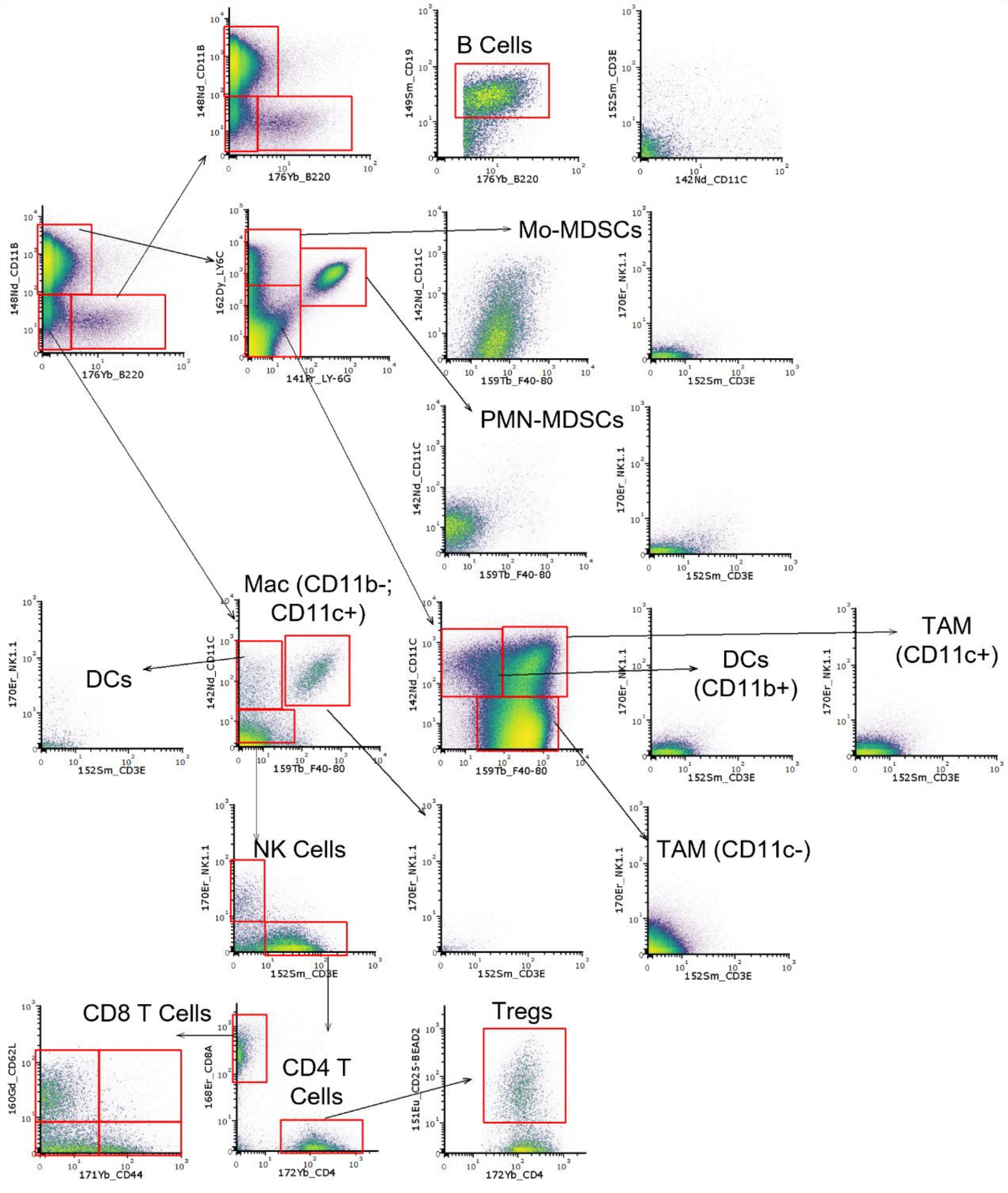


- CD11b- **Cd11c+** F4/80- → **DCs**
- CD11b- **Cd11c+ F4/80+** → **Mac (CD11b-; CD11c+)**
- **CD11b+ Cd11c+** F4/80- → **DCs (CD11b+)**
- **CD11b+ Cd11c+ F4/80+** → **TAM (CD11c+)**
- **CD11b+** Cd11c- **F4/80+** → **TAM (CD11c-)**

**EpCAM+**:

- **Trop2+ Ly6D+** CD104-: **Luminal-progenitor-like**
- **Trop2+** Ly6D- **CD104+**: **Progenitor-like**
- **Trop2+** Ly6D- CD104-: **Luminal-like**
- Trop2- Ly6D- **CD104+**: **Basal-like**

EpCAM-:

- **CD44+** CD104- **Ly6D+**: **EMT-like (Ly6D+)**
- **CD44+ CD104+** Ly6D-: **EMT-like (CD104+)**
- **CD44+** CD104- Ly6D-: **EMT-like**
- CD44- CD104- **Ly6D+**: **Ly6D+**
- CD44- **CD104+** Ly6D-: **CD104+**
- CD44- CD104- Ly6D-: **Other Stroma**

**Supplementary Figure 6. CyTOF Immune gating strategy.**

Gating strategy for Cd45$^{+}$ immune cells acquired from all individual tumours (n = 48) and analysed using a 23-antibody CyTOF panel combined with mass-tag barcoding. Specific nomenclature of immune and CD45$^{-}$ populations is indicated at the bottom.

### Supp Fig. 7

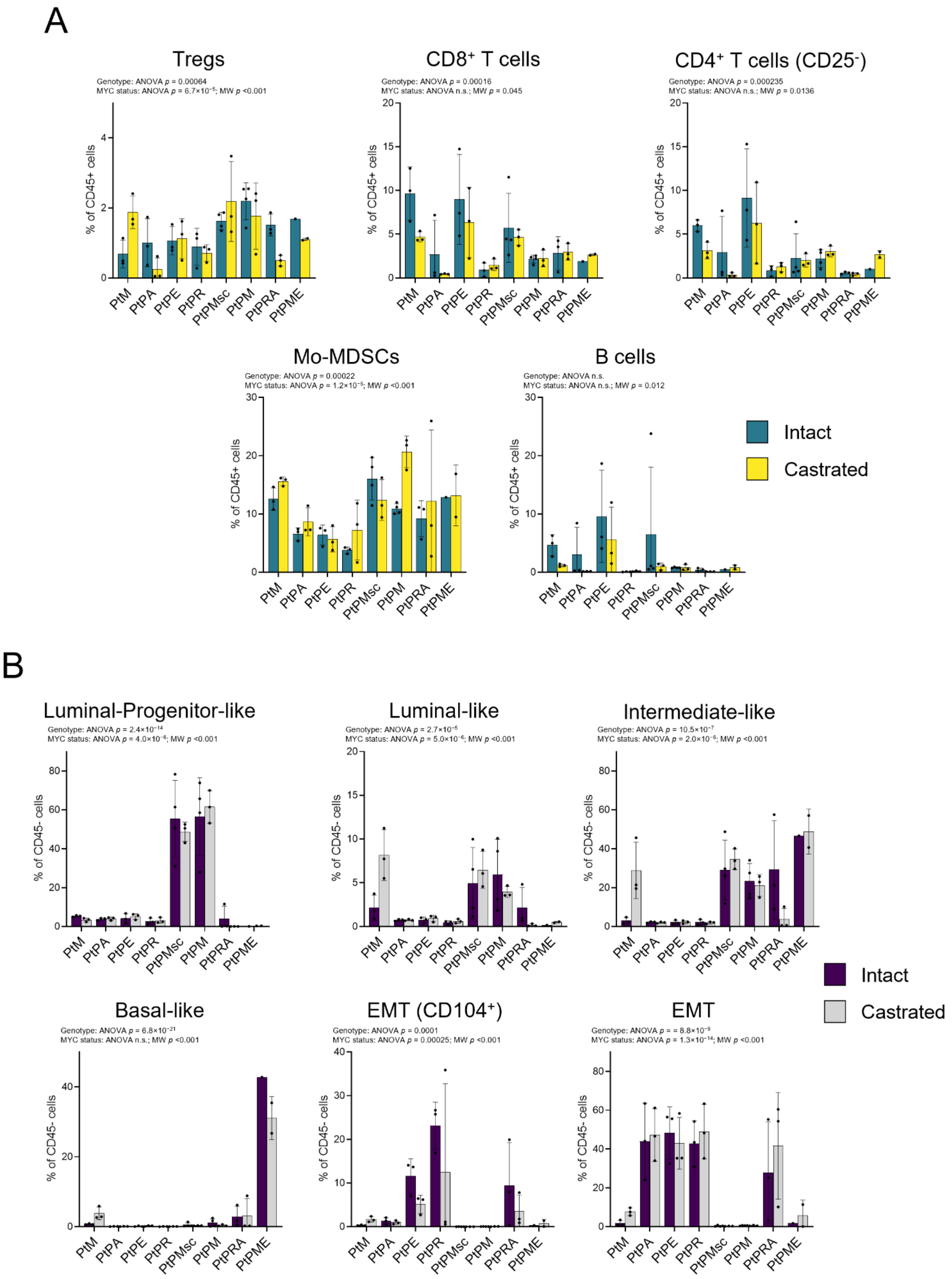

A
Tregs
Genotype: ANOVA p = 0.00064
MYC status: ANOVA p = 6.7×10⁻⁵; MW p <0.001
% of CD45+ cells
CD8⁺ T cells
Genotype: ANOVA p = 0.00016
MYC status: ANOVA n.s.; MW p = 0.045
% of CD45+ cells
CD4⁺ T cells (CD25⁻)
Genotype: ANOVA p = 0.000235
MYC status: ANOVA n.s.; MW p = 0.0136
% of CD45+ cells
Mo-MDSCs
Genotype: ANOVA p = 0.00022
MYC status: ANOVA p = 1.2×10⁻⁶; MW p <0.001
% of CD45+ cells
B cells
Genotype: ANOVA n.s.
MYC status: ANOVA n.s.; MW p = 0.012
% of CD45+ cells
PtM
PtPA
PtPE
PtPR
PtPMsc
PtPM
PtPRA
PtPME
Intact
Castrated
B
Luminal-Progenitor-like
Genotype: ANOVA p = 2.4×10⁻¹⁴
MYC status: ANOVA p = 4.0×10⁻⁶; MW p <0.001
% of CD45- cells
Luminal-like
Genotype: ANOVA p = 2.7×10⁻⁵
MYC status: ANOVA p = 5.0×10⁻⁶; MW p <0.001
% of CD45- cells
Intermediate-like
Genotype: ANOVA p = 10.5×10⁻⁷
MYC status: ANOVA p = 2.0×10⁻⁶; MW p <0.001
% of CD45- cells
Basal-like
Genotype: ANOVA p = 6.8×10⁻²¹
MYC status: ANOVA n.s.; MW p <0.001
% of CD45- cells
EMT (CD104⁺)
Genotype: ANOVA p = 0.0001
MYC status: ANOVA p = 0.00025; MW p <0.001
% of CD45- cells
EMT
Genotype: ANOVA p = = 8.8×10⁻⁹
MYC status: ANOVA p = 1.3×10⁻¹⁴; MW p <0.001
% of CD45- cells
Intact
Castrated

**Supplementary Figure 7. ProMPt tumours demonstrate genotype-specific immune and epithelial composition.**

(A) Bar plot showing replicate values for the indicated immune populations from CyTOF analysis. Two-way ANOVA was used to assess main and interaction effects of MYC status and castration. When overall effects were significant, pairwise Mann–Whitney U tests were performed to compare *MYC*$^+$ and *MYC*$^−$ groups within each condition. Bars represent mean ± SD (n = 3 per group). See also Figures 4 and S7B for extended results.

(B) Bar plot showing replicate values for the indicated Cd45$^-$ populations from CyTOF analysis. Two-way ANOVA was used to assess main and interaction effects of *MYC* status and castration. When overall effects were significant, pairwise Mann–Whitney U tests were performed to compare *MYC*$^+$ and *MYC*$^−$ groups within each condition. Bars represent mean ± SD (n = 3 per group). See also Figures 4 and S7A for extended results.

Supp Fig. 8

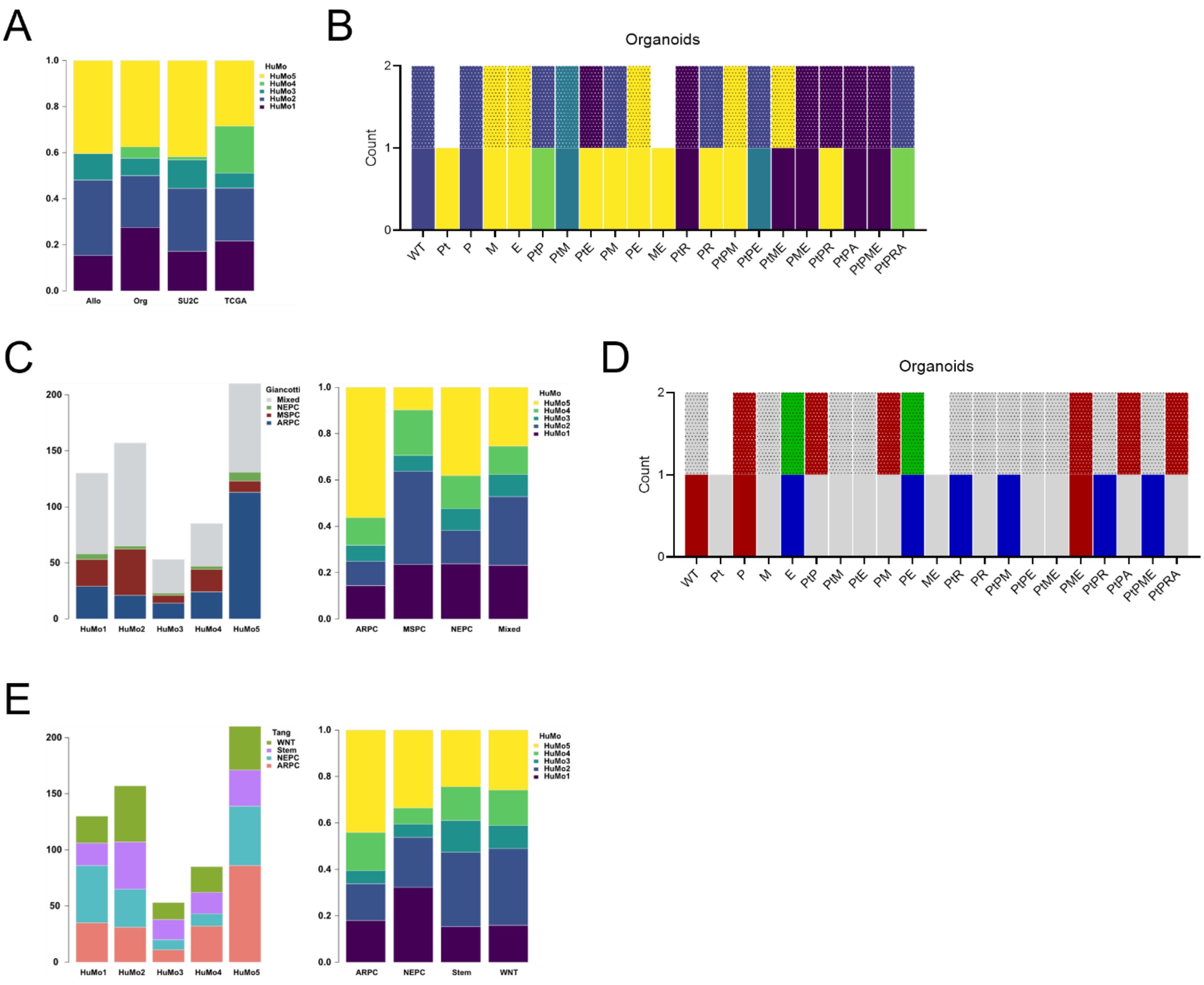


**Supplementary Figure 8. Pathway and subtype analysis of integrated human and mouse data highlights cluster specific characteristics.**

(A) Relative distribution of samples (Org: ProMPt organoids; Allo: Generation 2 ProMPt models) within individual HuMo clusters. Distributions are normalised within each group (sum = 1).

(B) ProMPt organoids classified by HuMo cluster, coloured by cluster identity. Patterned colours indicate organoids cultured in -DHT long condition.

(C) Distribution of samples coloured by subtype (ARPC, MSPC, NEPC and Mixed(10)) within individual HuMo clusters (left). Relative distribution of ARPC, MSPC, NEPC and Mixed(10) samples within individual HuMo clusters. Distributions are normalised within each group (sum = 1) (right).

(D) ProMPt organoids classified into ARPC, MSPC, NEPC, and Mixed subtypes using gene signatures from Han et al.(10). Patterned colours indicate organoids cultured in -DHT long condition.

(E) Distribution of samples coloured by subtype (ARPC, NEPC, STEM and WNT(8)) within individual HuMo clusters (left). Relative distribution of ARPC, NEPC, STEM and WNT(8) samples within individual HuMo clusters. Distributions are normalised within each group (sum = 1) (right).

Supp Fig. 9

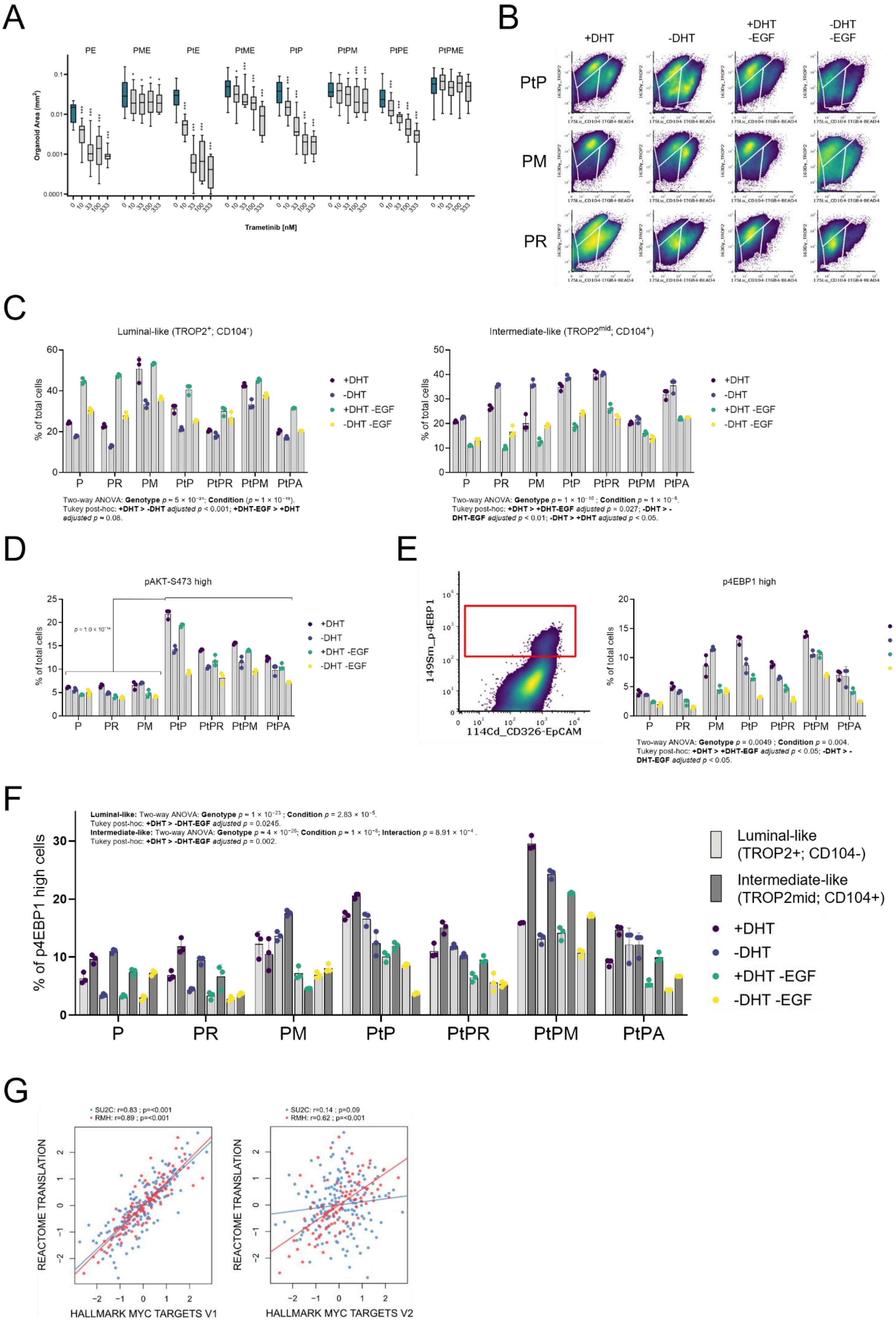
A
PE
PME
PtE
PtME
PtP
PtPM
PtPE
PtPME
Organoid Area (mm²)
Trametinib [nM]
B
+DHT
-DHT
+DHT -EGF
-DHT -EGF
PtP
PM
PR
C
Luminal-like (TROP2+; CD104-)
Intermediate-like (TROP2mid; CD104+)
% of total cells
+DHT
-DHT
+DHT -EGF
-DHT -EGF
P
PR
PM
PtP
PtPR
PtPM
PtPA
Two-way ANOVA: Genotype p ≈ 5 × 10⁻²¹; Condition (p ≈ 1 × 10⁻¹⁹).
Tukey post-hoc: +DHT > -DHT adjusted p < 0.001; +DHT-EGF > +DHT adjusted p ≈ 0.08.
Two-way ANOVA: Genotype p ≈ 1 × 10⁻¹⁰ ; Condition p ≈ 1 × 10⁻⁶.
Tukey post-hoc: +DHT > +DHT-EGF adjusted p = 0.027; -DHT > -DHT-EGF adjusted p < 0.01; -DHT > +DHT adjusted p < 0.05.
D
pAKT-S473 high
p = 1.0 × 10⁻¹⁴
E
149Sm_p4EBP1
114Cd_CD326-EpCAM
p4EBP1 high
Two-way ANOVA: Genotype p = 0.0049 ; Condition p = 0.004.
Tukey post-hoc: +DHT > +DHT-EGF adjusted p < 0.05; -DHT > -DHT-EGF adjusted p < 0.05.
F
Luminal-like: Two-way ANOVA: Genotype p ≈ 1 × 10⁻²³ ; Condition p = 2.83 × 10⁻⁵.
Tukey post-hoc: +DHT > -DHT-EGF adjusted p = 0.0245.
Intermediate-like: Two-way ANOVA: Genotype p ≈ 4 × 10⁻²⁶; Condition p ≈ 1 × 10⁻⁸; Interaction p = 8.91 × 10⁻⁴ .
Tukey post-hoc: +DHT > -DHT-EGF adjusted p = 0.002.
% of p4EBP1 high cells
Luminal-like (TROP2+; CD104-)
Intermediate-like (TROP2mid; CD104+)
G
SU2C: r=0.83 ; p=<0.001
RMH: r=0.89 ; p=<0.001
SU2C: r=0.14 ; p=0.09
RMH: r=0.62 ; p=<0.001
REACTOME TRANSLATION
HALLMARK MYC TARGETS V1
HALLMARK MYC TARGETS V2

**Supplementary Figure 9. Characterization of epithelial populations following media permutation.**

(A) MYC-driven resistance to MAPK inhibition assessed by changes in organoid area ($mm^2$; n = 50) following increasing doses of trametinib (nM). *$P < 0.05$; **$P < 0.01$; ***$P < 0.001$ by Mann–Whitney U test compared with the untreated control for each genotype.

(B) Gated density plots showing luminal-like (Trop2$^{+}$; CD104$^{-}$) and intermediate-like (Trop2$^{mid}$; CD104$^{+}$) populations across selected ProMPt genotypes. Data represent CyTOF media permutation conditions (+DHT, –DHT, +DHT–EGF, –DHT–EGF) corresponding to Figure 6B.

(C) Percentage of luminal-like (left) and intermediate-like (right) cells under control, –DHT, and EGF-deprived conditions in the CyTOF media permutation assay. Two-way ANOVA was used to assess main and interaction effects of genotype and culture condition. When overall effects were significant, Tukey's post hoc test was performed for pairwise comparisons. Significant comparisons are indicated below the plots. Non-parametric Mann–Whitney U tests were applied within genotypes to confirm specific differences. Bars represent mean ± SD (n = 3 per group).

(D) Percentage of pAKT$^{high}$ cells in the CyTOF media permutation assay. Two-tailed Mann–Whitney U test was used to assess differences between *Pten*-intact and *Pten*-deficient genotypes. Bars represent mean ± SD (n = 3 per group).

(E) Density plot (left) visualizing the gating of the p4EBP1 high cells. Analysis of p4EBP1$^{high}$ cells in the CyTOF media permutation assay (right). Two-way ANOVA was used to assess main and interaction effects of genotype and culture condition. When overall effects were significant, Tukey's post hoc test was performed for pairwise comparisons. Significant comparisons are indicated below the plots. Non-parametric Mann–Whitney U tests were applied within genotypes to confirm specific differences. Bars represent mean ± SD (n = 3 per group).

(F) Percentage of p4EBP1$^{high}$ cells within luminal-like and intermediate-like populations in the CyTOF media permutation assay. Two-way ANOVA was used to assess main and interaction effects of genotype and culture condition. When overall effects were significant, Tukey's post hoc test was applied for pairwise comparisons. Significant comparisons are indicated below the plots. Non-parametric Mann–Whitney U tests were used within genotypes to confirm specific differences. PtPM retained the highest fraction of p4EBP1$^{high}$ cells across all genotypes in both luminal and intermediate compartments. In intermediate cells, PtPM showed significantly higher mean levels than all other genotypes ($P < 0.001$ for all pairwise comparisons). Across treatment conditions, intermediate cells in the PtPM genotype consistently showed higher p4EBP1$^{high}$ levels than luminal cells (P = 0.0006). Bars represent mean ± SD (n = 3 per group).

(G) Scatterplots showing correlations between Hallmark MYC Targets (V1, left; V2, right) and the Reactome Translation signature across the SU2C and Royal Marsden Hospital datasets. Linear fits were calculated per dataset, with corresponding Pearson correlation coefficients and *P* values indicated.

Supp Fig. 10

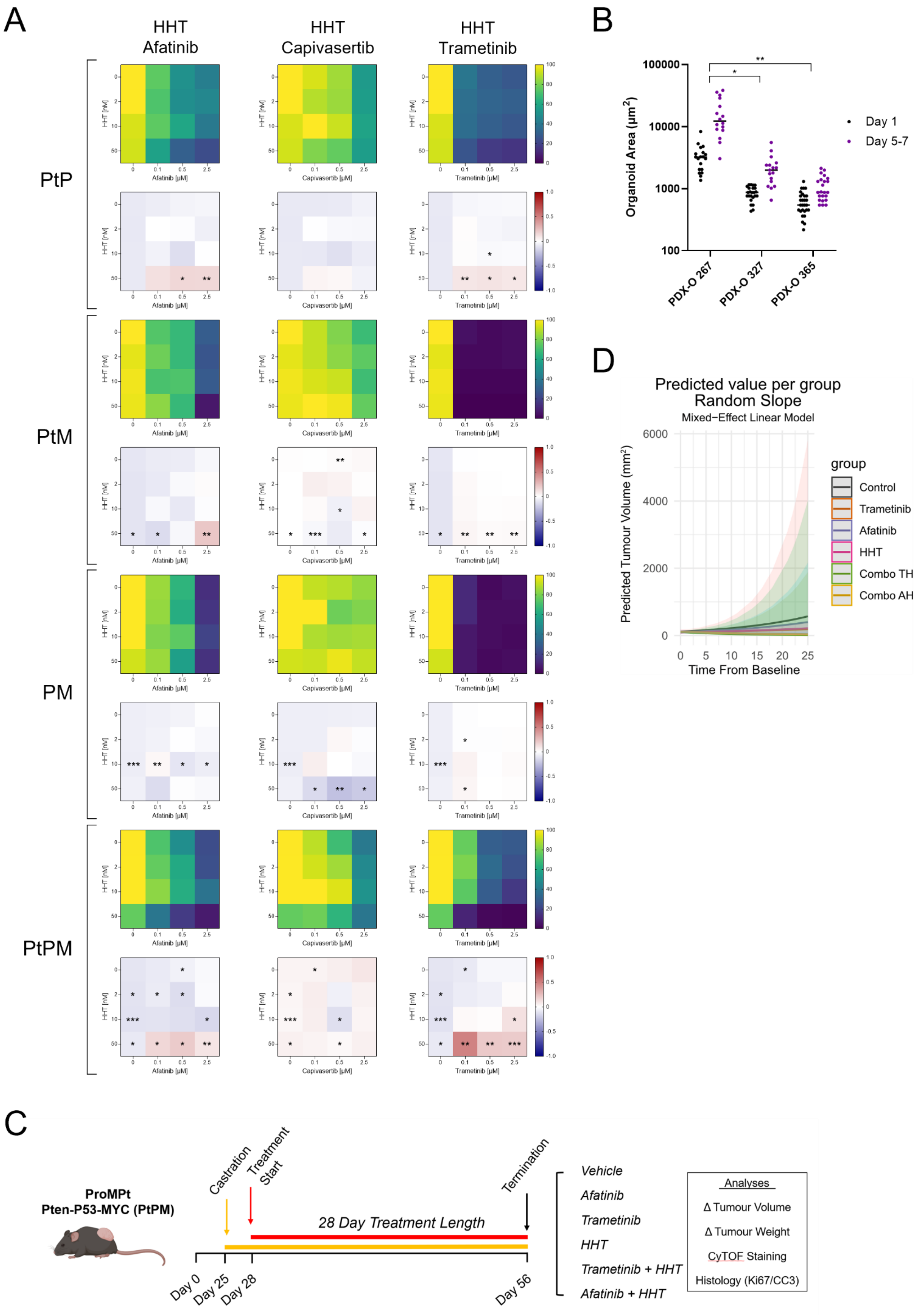
A
HHT
Afatinib
HHT
Capivasertib
HHT
Trametinib
PtP
PtM
PM
PtPM
HHT [nM]
Afatinib [µM]
Capivasertib [µM]
Trametinib [µM]
B
Organoid Area (µm²)
100000
10000
1000
100
PDX-O 267
PDX-O 327
PDX-O 365
Day 1
Day 5-7
D
Predicted value per group
Random Slope
Mixed−Effect Linear Model
Predicted Tumour Volume (mm³)
6000
4000
2000
0
0
5
10
15
20
25
Time From Baseline
group
Control
Trametinib
Afatinib
HHT
Combo TH
Combo AH
C
ProMPt
Pten-P53-MYC (PtPM)
Castration
Treatment
Start
Termination
28 Day Treatment Length
Day 0
Day 25
Day 28
Day 56
Vehicle
Afatinib
Trametinib
HHT
Trametinib + HHT
Afatinib + HHT
Analyses
Δ Tumour Volume
Δ Tumour Weight
CyTOF Staining
Histology (Ki67/CC3)

**Supplementary Figure 10. Inhibition of MAPK and Protein Translation synergize to inhibit tumour development**.

(A) Viability heatmap (top) and Bliss synergy score map (bottom) for ProMPt organoids treated with Afatinib, Capivasertib or Trametinib in combination with HHT. A two-sided one-sample t-test was applied to triplicate inhibition fractions at each dose pair to assess deviation from Bliss additivity. Asterisks denote dose pairs where observed inhibition significantly exceeded the Bliss expectation (*$P < 0.05$, **$P < 0.01$, ***$P < 0.001$).

(B) Growth analysis of three PDX-derived organoid (PDX-O) populations based on area measurements at day 1 and endpoint (day 5 for PDX-O 267; day 7 for the other two). Growth rates were calculated as the ratio of organoid area (endpoint/day 1) and compared using Welch's t-test (*$P < 0.05$, **$P < 0.01$).

(C) Timeline of the *in vivo* ProMPt preclinical trial in C57BL/6 mice. Subcutaneous PtPM organoids ($1.5 \times 10^6$) were implanted and allowed to develop for 25 days before castration and treatment initiation.

(D) Predicted tumour growth of PtPM therapeutic cohorts, modelled using a linear mixed-effects regression.

Supp Fig. 11

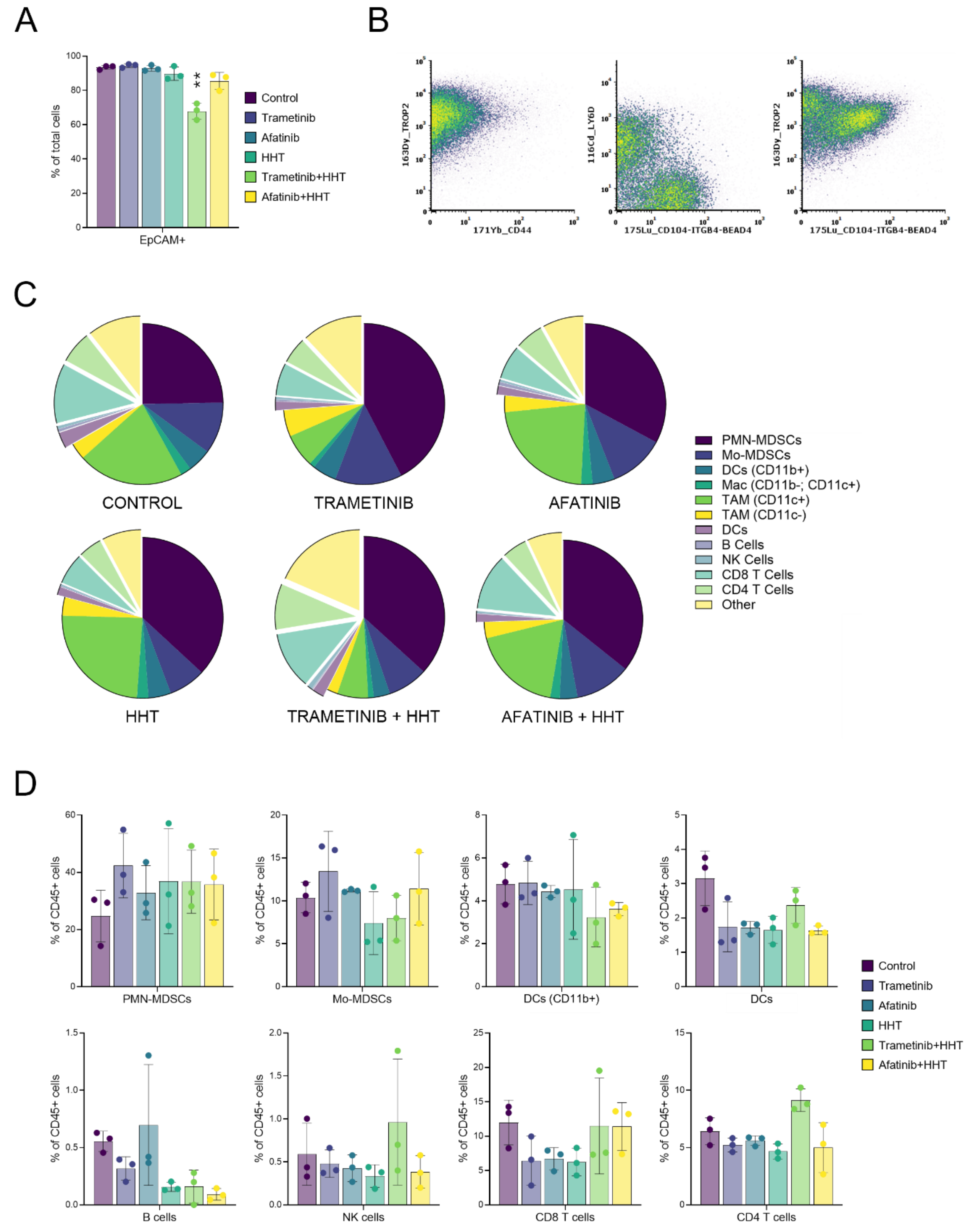


**Supplementary Figure 11. Therapeutic modulation of immune populations.**

(A) Abundance of EpCAM+ in endpoint PtPM tumours across treatment arms, quantified by CyTOF analysis. Comparisons with vehicle controls were performed using Welch's t-test (***P* < 0.01). Bars indicate mean ± SD (n = 3 per group).

(B) Density plot showing EpCAM+ cells from pooled CyTOF data of the PtPM treatment experiment. All cells show Trop2 positivity (left) and are separated into distinct populations by Ly6D and CD104 (middle). CD104+ cells also show positive TROP2 levels (right).

(C) Total immune populations detected via CyTOF from treated PtPM tumours. Data generated from 3 biological replicates per cohort.

(D) Relative abundance of the indicated cell populations in endpoint PtPM tumours across treatment arms. Bars indicate mean ± SD (n = 3 per group).
