## Supplementary material for "ProMPt: A modular preclinical platform for functional modelling of prostate cancer heterogeneity and therapeutic vulnerabilities": Document S2 - Supp tables 4-5

**Table S4**. Genomic details of PDX-O samples. Related to Figure 7 and S10.

| ID | MYC | P53 | PTEN | RB1 | ERG | AR | Genomics |
|---|---|---|---|---|---|---|---|
| CP327 | High | Neg/Mut | Positive | Positive | Negative | Positive | ARID2 p.C382Y; p.T426M; FANCD2 p. F925Sfs*17; FANCF p. L162Dfs*103; TP53 p. R196 |
| CP267 | Positive | Neg/Mut | Negative | Negative | Negative | Negative | AR p.L702H; AR p.S889G; PIK3CG p.R684L; TP53. R248Q |
| CP365 | Negative | Negative | Positive | Positive | Negative | Positive | CDKN1B, RB1, WT1 deep deletions; BRCA1 amplification |

**Table S4** Status of c-MYC, TP53, PTEN, RB1, ERG, and AR expression assessed through IHC as previously described (Guo *et al*. 2023). Results of targeted next-generation sequencing (CP327) or whole genome sequencing provided.

**Table S5**. CyTOF Antibody Panels. Related Figures 4, 6, 7, and Figure S6, S7, S9, and S11.

***Organoids media permutation panel***

| Target | Clone | Metal | Supplier |
|---|---|---|---|
| Phospho-Histone H3 [Ser28] | HTA28 | 89Y | Standard BioTools |
| Cytokeratin 8 | EP1628Y | 113Cd | Abcam |
| CD326/EpCAM | G8.8 | 114Cd | BioLegend |
| Clusterin | Poly | 115In | Bio-Techne |
| Ly-6D | 49-H4 | 116Cd | BD Biosciences |
| TOBis 126-Plex | - | 122Te | |
| TOBis 126-Plex | - | 123Te | |
| TOBis 126-Plex | - | 124Te | |
| TOBis 126-Plex | - | 125Te | |
| TOBis 126-Plex | - | 126Te | |
| IdU (S-Phase) | - | 127I | |
| TOBis 126-Plex | - | 128Te | |
| TOBis 126-Plex | - | 130Te | |
| CD49f | GoH3 | 141Pr | BioLegend |
| Cleaved Caspase 3 | D3E9 | 142Nd | Standard BioTools |
| Phospho-SEK1/MKK4 [Ser257] | C36C11 | 143Nd | CST |
| CD44 | IM7 | 144Nd | BioLegend |
| Phospho-BTK [Tyr551] | Y511 | 147Sm | BD Biosciences |
| Phospho-p90RSK [Thr359] | D1E9 | 148Sm | CST |
| Phospho-4E-BP1 [Thr37/46] | 236B4 | 149Sm | Standard BioTools |
| Phospho-AMPKα [Thr172] | 40H9 | 151Eu | CST |
| Phospho-Akt [Ser473] | DE9 | 152Sm | Standard BioTools |
| Phospho-GSK-3β [Ser9] | D85E12 | 153Eu | CST |
| Phospho-NDRG1 [Thr346] | D98G11 | 154Sm | CST |
| Phospho-38 [Thr180/Tyr182] | D3F9 | 156Gd | Standard BioTools |
| Phospho-SRC [Tyr418] | SC1T2M3 | 157Gd | Thermo Fisher |
| Phospho-FAK [Tyr397] | Poly | 158Gd | CST |
| Phospho-MAPKAPK2 [Thr334] | 27B7 | 159Tb | Standard BioTools |
| CD24 | M1/69 | 160Gd | BioLegend |
| Phospho-Bad [Ser112] | 40A9 | 161Dy | Standard BioTools |
| TROP2 | Poly | 163Dy | Bio-Techne |
| CyclinB1 | GNS-1 | 164Dy | Standard BioTools |
| Phospho-MEK1/2 [Ser221] | 166F8 | 165Ho | CST |
| Phospho-NFkBp65 [S529] | K10x | 166Er | Standard BioTools |
| Phospho-ERK1/2 [Thr202/Tyr204] | D1314.4E | 167Er | Standard BioTools |
| Phospho-SMAD2 [Ser465/467]/ SMAD3 [Ser423/425] | D27F4 | 168Er | CST |
| Ly-6A/E /Sca-1 | D7 | 169Tm | Standard BioTools |

| | | | |
|---|---|---|---|
| Phospho-Akt [Thr308] | J1-223.317 | 171Yb | BD Biosciences |
| Phospho-S6 [Ser235/236] | N7-548 | 172Yb | Standard BioTools |
| CD133/Prom1 | 315-2C11 | 173Yb | BioLegend |
| CD104/ITGB4 | 346-11A | 175Lu | BioLegend |
| Phospho-CREB [Ser133] | 87G3 | 176Yb | Standard BioTools |
| DNA Intercalator ($^{191/193}$Ir) | - | 191/193Ir | Standard BioTools |
| Viability ($^{194}$Cisplatin) | - | 194Pt | Standard BioTools |
| TOBis 126-Plex | - | 196Pt | |
| TOBis 126-Plex | - | 198Pt | |

***Immune phenotyping panel***

| Target | Clone | Metal | Supplier |
|---|---|---|---|
| Cell-ID Barcode | - | 102Pd | Standard BioTools |
| Cell-ID Barcode | - | 104Pd | Standard BioTools |
| Cell-ID Barcode | - | 105Pd | Standard BioTools |
| Cell-ID Barcode | - | 106Pd | Standard BioTools |
| Cell-ID Barcode | - | 108Pd | Standard BioTools |
| Cell-ID Barcode | - | 110Pd | Standard BioTools |
| Ly-6D | 49-H4 | 116Cd | BD Biosciences |
| Ly-6G | 1A8 | 141Pr | Standard BioTools |
| CD11c (#201306) | N418 | 142Nd | Standard BioTools |
| CD69 (#201306) | H1.2F3 | 145Nd | Standard BioTools |
| CD45 (#201306) | 30-F11 | 147Sm | Standard BioTools |
| CD11b (#201306) | M1/70 | 148Nd | Standard BioTools |
| CD19 (#201306) | 6D5 | 149Sm | Standard BioTools |
| CD25 (#201306) | 3C7 | 151Eu | Standard BioTools |
| CD3e (#201306) | 145-2C11 | 152Sm | Standard BioTools |
| Ter-119 (#201306) | Ter-119 | 154Sm | Standard BioTools |
| F4/80 | BM8 | 159Tb | Standard BioTools |
| CD62L (#201306) | MEL-14 | 160Gd | Standard BioTools |
| Ly-6C | HK1.4 | 162Dy | Standard BioTools |
| TROP2 | - | 163Dy | Bio-Techne |
| CD326/EpCAM | G8.8 | 166Er | Standard BioTools |
| CD8a (#201306) | 53-6.7 | 168Er | Standard BioTools |
| CD206/MMR | C068C2 | 169Tm | Standard BioTools |
| NK1.1 (#201306) | PK136 | 170Er | Standard BioTools |
| CD44 (#201306) | IM7 | 171Yb | Standard BioTools |
| CD4 (#201306) | RM4-5 | 172Yb | Standard BioTools |

| | | | |
|---|---|---|---|
| CD133/Prom1 | 315-2C11 | 173Yb | BioLegend |
| CD104/ITGB4 | 346-11A | 175Lu | BioLegend |
| B220 (#201306) | RA3-6B2 | 176Yb | Standard BioTools |
| DNA Intercalator ($^{191/193}$Ir) | - | 191/193Ir | Standard BioTools |
| Viability ($^{196/198}$Cisplatin) | - | 196198Pt | Standard BioTools |

**Table S5.** CyTOF antibody panels used across experimental conditions.
Immune phenotyping panel: Figure 4A-C, Figure S6-7, Figure 7E-G, and Figure S11A-D.
Organoids media permutation panel: Figure 6B-D, and Figure S9B-F.
Extracellular (green), intracellular (blue), non-antibody material (orange).
